# Phylogenomic analyses of echinoid diversification prompt a re-evaluation of their fossil record

**DOI:** 10.1101/2021.07.19.453013

**Authors:** Nicolás Mongiardino Koch, Jeffrey R Thompson, Avery S Hatch, Marina F McCowin, A Frances Armstrong, Simon E Coppard, Felipe Aguilera, Omri Bronstein, Andreas Kroh, Rich Mooi, Greg W Rouse

## Abstract

Echinoids are key components of modern marine ecosystems. Despite a remarkable fossil record, the emergence of their crown group is documented by few specimens of unclear affinities, rendering much of their early history uncertain. The origin of sand dollars, one of its most distinctive clades, is also unclear due to an unstable phylogenetic context and discrepancies between molecular divergence times and fossil evidence. We employ seventeen novel genomes and transcriptomes to build a phylogenomic dataset with a near-complete sampling of major lineages. With it, we revise the phylogeny and divergence times of echinoids, and place their history within the broader context of echinoderm evolution. We also introduce the concept of a chronospace—a multidimensional representation of node ages—and use it to explore the effects of using alternative gene samples, models of molecular evolution, and clock priors. We find the choice of clock model to have the strongest impact on divergence times, while the use of site-heterogeneous models shows little effects. The choice of loci shows an intermediate impact, affecting mostly deep Paleozoic nodes, for which clock-like genes recover dates more congruent with fossil evidence. Our results reveal that crown group echinoids originated in the Permian and diversified rapidly in the Triassic, despite the relative lack of fossil evidence for this early diversification. We also clarify the relationships among sand dollars and their close relatives, showing that the genus *Apatopygus* represents a relict lineage with a deep Jurassic origin. Surprisingly, the origin of sand dollars is confidently dated to the Cretaceous, implying ghost ranges spanning approximately 50 million years, a remarkable discrepancy with their rich fossil record.

## Introduction

The fossil record represents the best source of primary data for constraining the origins of major lineages across the tree of life. However, the fossil record is not perfect, and even for groups with an excellent fossilization potential, constraining their age of origin can be difficult [1, 2]. Furthermore, as many traditional hypotheses of relationships have been revised in light of large-scale molecular datasets, the affinities of fossil lineages and their bearings on inferred times of divergence have also required a reassessment. An exemplary case of this is Echinoidea, a clade comprised of sea urchins, heart urchins, sand dollars, and allies, for which phylogenomic trees have questioned the timing of previously well-constrained nodes [3, 4].

Echinoids are easily recognized by their spine-covered skeletons or tests, composed of numerous tightly interlocking plates. Slightly over 1,000 living species have been described to date [5], a diversity that populates every marine benthic environment from intertidal to abyssal depths [6].

Echinoids are usually subdivided into two morpho-functional groups with similar species-level diversities: “regular” sea urchins, a paraphyletic assemblage of hemispherical, epibenthic consumers protected by large spines; and irregulars (Irregularia), a clade of predominantly infaunal and bilaterally symmetrical forms covered by small and specialized spines. In today’s oceans, regular echinoids act as ecosystem engineers in biodiverse coastal communities such as coral reefs [7] and kelp forests [8], where they are often the main consumers. They are first well-known in the fossil record on either side of the Permian-Triassic (P-T) mass extinction event when many species occupied reef environments similar to those inhabited today by their descendants [9, 10]. This extinction event was originally thought to have radically impacted the macroevolutionary history of the clade, decimating the echinoid stem group and leading to the radiation of crown group taxa from a single surviving lineage [11, 12]. However, it is now widely accepted that the origin of crown group Echinoidea (i.e., the divergence between its two main lineages, Cidaroidea and Euechinoidea) occurred in the Late Permian, as supported by molecular estimates of divergence [13, 14], as well as the occurrence of Permian fossils with morphologies typical of modern cidaroids [15, 16]. However, a recent total-evidence study recovered many taxa previously classified as crown group members along the echinoid stem, while also suggesting that up to three crown group lineages survived the P-T mass extinction [3]. This result increases the discrepancy between molecular estimates and the fossil record and renders uncertain the early evolutionary history of crown group echinoids. Constraining the timing of origin of this clade relative to the P-T mass extinction (especially in the light of recent topological changes [3, 4]) is further complicated by the poor preservation potential of stem group echinoids, and the difficulty assigning available disarticulated remains from the Late Paleozoic and Early Triassic to specific clades [11, 12, 17–20].

Compared to the morphological conservatism of regular sea urchins, the evolutionary history of the relatively younger Irregularia was characterized by dramatic levels of morphological and ecological innovation [21–24]. Within the diversity of irregulars, sand dollars are the most easily recognized (Fig. 1). The clade includes greatly flattened forms that live in high-energy sandy environments where they feed using a unique mechanism for selecting and transporting organic particles to the mouth, where these are crushed using well-developed jaws [25, 26]. Sand dollars (Scutelloida) were long thought to be most closely related to sea biscuits (Clypeasteroida) given a wealth of shared morphological characters [19, 25]. The extraordinary fossil record of both sand dollars and sea biscuits suggested their last common ancestor originated in the early Cenozoic from among an assemblage known as “cassiduloids” [23, 25], a once diverse group that is today represented by three depauperate lineages: cassidulids (and close relatives), echinolampadids, and apatopygids [19, 27]. These taxa not only lack the defining features of both scutelloids and clypeasteroids but have experienced little morphological change since their origin deep in the Mesozoic [24, 27–29]. However, early molecular phylogenies supported both cassidulids and echinolampadids as close relatives of sand dollars (e.g., [14, 30]), a topology initially disregarded for its conflicts with both morphological and paleontological evidence, but later confirmed using phylogenomic approaches [4]. While many of the traits shared by sand dollars and sea biscuits have since been suggested to represent a mix of convergences and ancestral synapomorphies secondarily lost by some “cassiduloids” [3, 4], the strong discrepancy between molecular topologies and the fossil record remains unexplained. Central to this discussion is the position of apatopygids, a clade so far unsampled in molecular studies. Apatopygids have a fossil record stretching more than 100 million years and likely have phylogenetic affinities with even older extinct lineages [3, 19, 28, 29]. Although current molecular topologies already imply ghost ranges for scutelloids and clypeasteroids that necessarily extend beyond the Cretaceous-Paleogene (K-Pg) boundary, the phylogenetic position of apatopygids could impose even earlier ages on these lineages (Fig. 1). Constraining these divergences is necessary to understand the timing of origin of the sand dollars, one of the most specialized lineages of echinoids [24–27]. Resolving some phylogenetic relationships within scutelloids has also been complicated by their recurrent miniaturization and associated loss of morphological features (Fig. 1; [25, 31, 32]).

**Figure 1:**
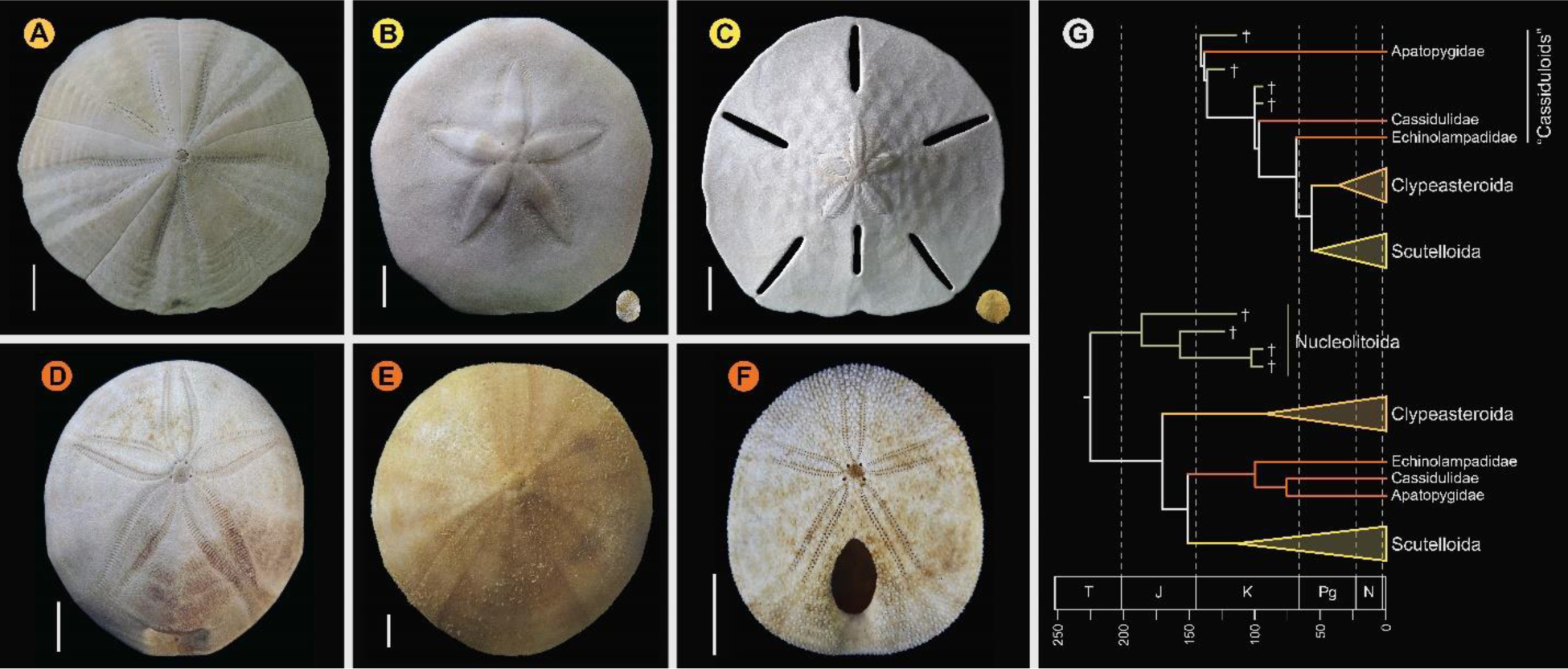
Neognathostomate diversity and phylogenetic relationships. **A.** *Fellaster zelandiae*, North Island, New Zealand (Clypeasteroida). **B.** Large specimen: *Peronella japonica*, Ryukyu Islands, Japan; Small specimen: *Echinocyamus crispus*, Maricaban Island, Philippines (Laganina: Scutelloida). **C.** Large specimen: *Leodia sexiesperforata*, Long Key, Florida; Small specimen: *Sinaechinocyamus mai*, Taiwan (Scutellina: Scutelloida). **D.** *Rhyncholampas pacificus,* Isla Isabela, Galápagos Islands (Cassidulidae). **E.** *Conolampas sigsbei*, Bimini, Bahamas (Echinolampadidae). **F.** *Apatopygus recens*, Australia (Apatopygidae). **G.** Hypotheses of relationships among neognathostomates. Top: Morphology supports a clade of Clypeasteroida + Scutelloida originating after the K-Pg boundary, subtended by a paraphyletic assemblage of extant (red) and extinct (green) “cassiduloids” [19]. Bottom: A recent total-evidence study split most of cassiduloid diversity into a clade of extant lineages closely related to scutelloids, and an unrelated clade of extinct forms (Nucleolitoida) [3]. Divergence times are much older and conflict with fossil evidence. Cassidulids and apatopygids lacked molecular data in this analysis. Scale bars = 10 mm.

Echinoidea constitutes a model clade in developmental biology and genomics. As these fields embrace a more comparative approach [13, 33, 34], robust and time-calibrated phylogenies are expected to play an increasingly important role. Likewise, the extraordinary fossil record of echinoids and the ease with which echinoid fossils can be incorporated in phylogenetic analyses make them an ideal system to explore macroevolutionary dynamics using phylogenetic comparative methods [3, 31]. In this study, we build upon available molecular resources with 17 novel genome-scale datasets and build the largest molecular matrix for echinoids yet compiled. Our expanded phylogenomic dataset extends sampling to 16 of the 17 currently recognized echinoid orders (plus the unassigned apatopygids) [35], and is the first to bracket the extant diversity of both sand dollars and sea biscuits and include members of all three lineages of living “cassiduloids” (cassidulids, echinolampadids, and apatopygids). We also incorporate a diverse sample of outgroups, providing access to the deepest nodes within the crown groups of all other echinoderm classes (holothuroids, asteroids, ophiuroids, and crinoids). With it, we reconstruct the phylogenetic relationships and divergence times of the major lineages of living echinoids and place their diversification within the broader context of echinoderm evolution.

## Results

### **(a)** Phylogeny of Echinoidea

Phylogenetic relationships supported by the full dataset were remarkably stable, with all nodes but one being identically resolved and fully supported across a wide range of inference methods (Fig. 2A). While recovering a topology similar to those of previous molecular studies [3, 4, 13, 14, 30, 36], this analysis is the first to sample and confidently place micropygoids and aspidodiadematoids within Aulodonta, as well as resolve the relationships among all major clades of Neognathostomata (scutelloids, clypeateroids and the three lineages of extant “cassiduloids”). Our results show that *Apatopygus recens* is not related to the remaining “cassiduloids” but is instead the sister clade to all other sampled neognathostomates. The strong support for this placement, as well as for a clade of cassidulids and echinolampadids (Cassiduloida *sensu stricto*) as the sister group to sand dollars, provides a basis for an otherwise elusive phylogenetic classification of neognathostomates. Our topology also confirms that *Sinaechinocyamus mai*, a miniaturized species once considered a plesiomorphic member of Scutelloida based on the reduction or loss of diagnostic features (Fig. 1), is in fact a derived paedomorphic lineage closely related to *Scaphechinus mirabilis* [32].

**Figure 2:**
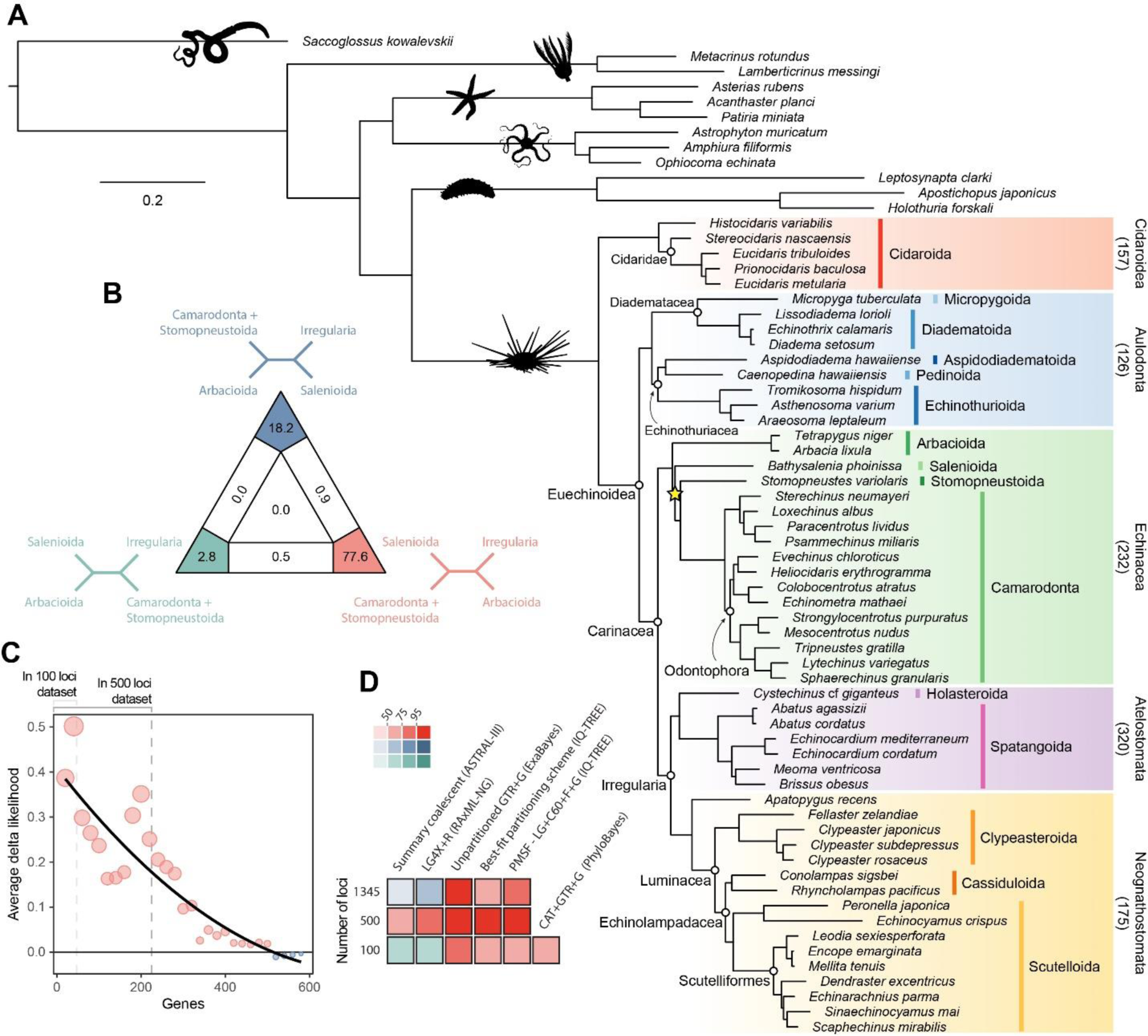
Phylogenetic relationships among major clades of Echinoidea. **A.** Favored topology. With the exception of a single contentious node within Echinacea (marked with a star), all methods supported the same pattern of relationships. Numbers below major clades correspond to the current numbers of described living species (obtained from [5]). **B.** Likelihood-mapping analysis showing the proportion of quartets supporting different resolutions within Echinacea. While the majority of quartets support the topology depicted in **A** (shown in red), a significant number support an alternative resolution that has been recovered in morphological analyses [19] (shown in blue). **C.** Support for a clade of Salenioida + (Camarodonta + Stomopneustoida), as depicted in **A**, is predominantly concentrated among the most phylogenetically useful loci. This signal is attenuated in larger datasets that contain less reliable genes, eventually favoring an alternative resolution. Only the 584 loci containing data for the three main lineages of Echinacea were considered. The line corresponds to a second-degree polynomial regression. **D.** Resolution and bootstrap scores (color coded) of the topology within Echinacea found using datasets of different sizes and alternative methods of inference.

Salenioida is another major lineage sampled here for the first time, and whose exact position among regular echinoids proved difficult to resolve. While some methods supported salenioids as the sister group to a clade of camarodonts, stomopneustoids, and arbacioids (a topology previously supported by morphology [19]), others recovered a closer relationship of salenioids to Camarodonta + Stomopneustoida, with arbacioids sister to them all (as shown in Fig. 2A). As revealed using likelihood mapping, these results do not stem from a lack of phylogenetic signal, but rather from the presence of strong and conflicting evidence in the dataset regarding the position of salenioids (Fig. 2B). However, a careful dissection of these signals shows that loci with high phylogenetic usefulness (as defined by [3, 37]; see Methods) favor the topology shown in Fig. 2A, with the morphological hypothesis becoming dominant only after incorporating less reliable loci (Fig .2C). In line with these results, moderate levels of gene subsampling (down to 500 loci) targeting the most phylogenetically useful loci unambiguously support the placement of arbacioids as sister to the remaining taxa, regardless of the chosen method of inference (Fig. 2D). More extreme subsampling (down to 100 loci) again results in disagreement among methods. This possibly stems from the increasing effect of stochastic errors in smaller datasets, as less than half of the sampled loci in these reduced datasets contain data for all branches of this quartet (see Fig. 2C). This result shows the importance of ensuring that datasets (especially subsampled ones) retain appropriate levels of occupancy for clades bracketing contentious nodes [38]. Despite these disagreements, several lines of evidence favor the topology shown in Fig. 2A, including the results of likelihood mapping, and the increased support for this resolution among the most phylogenetically useful loci and when using more complex methods of reconstruction, such as partitioned and site-heterogeneous models, which always favor this topology regardless of dataset size (Fig. 2D).

### **(b)** Sensitivity of node ages

While alternative methods of inference had minor effects on phylogenetic relationships, they did impact the reconstruction of branch lengths (Fig. S1). Site-heterogeneous models (such as CAT+GTR+G) returned longer branch lengths overall, but also uncovered a larger degree of molecular change among echinoderm classes. Branches connecting these clades were stretched to a much larger extent than those within the ingroup (Fig. S1), which might affect the inference of node dates. We tested this hypothesis by exploring the sensitivity of divergence times to the use of alternative models of molecular evolution (site-homogeneous vs. site-heterogeneous), as well as different clock priors (autocorrelated vs. uncorrelated), and gene sampling strategies (using five different approaches; see Methods). All combinations of these factors were explored, resulting in 20 different time-calibration settings that were run using Bayesian approaches under a constrained tree topology (that shown in Fig. 2A). Posterior distributions of node ages were then obtained from each of these time-calibrated analyses, confirming that inferred divergence times were significantly affected by all of these factors (multivariate analysis of variance; all *P* < 2.2x10^-16^). While nodes connecting some outgroup taxa were among those most sensitive to these decisions, large effects were also seen among nodes relating to the origin and diversification of the echinoid clades Cidaroidea and Aulodonta. All of these nodes varied in age by more than 60 Ma—and up to 115 Ma—among the consensus topologies of different analyses (Fig. S2).

In order to isolate and visualize the impact of each of these factors on divergence time estimation, chronograms were represented in a multidimensional space of node dates, with each axis representing the age of a given node. We term this type of graph a chronospace given its similarities to the treespaces commonly used to explore topological differences between phylogenetic trees [39]. Each observation (chronogram) was classified as obtained under a specific clock prior, model of molecular evolution, and gene sampling strategy, and the major effects of each of these choices were extracted with the use of between-group principal component analyses (bgPCAs). This method of ordination allows for the isolation of axes that capture differences between group means, and has been used before with other high-dimensional datasets such as obtained from microarrays and geometric morphometric approaches [40, 41]. The single dimension of chronospace maximizing the distinctiveness of chronograms obtained under different clock models explained 53.7% of the total variance in node ages across all analyses (Fig. 3). In contrast, the choice of different loci or models of molecular evolution showed a much lesser effect, explaining only 10.5% and 4.0% of the total variance, respectively. Even though all of these decisions affected a similar set of sensitive nodes (those mentioned above, as well as some relationships within Atelostomata), the choice of clock model modified the ages of 17 of these by more than 20 Ma (Fig. S3). This degree of change was induced on only 5 nodes by selecting alternative loci (Fig. S4), and was not induced on any node by enforcing different models of evolution (Fig. S5).

**Figure 3:**
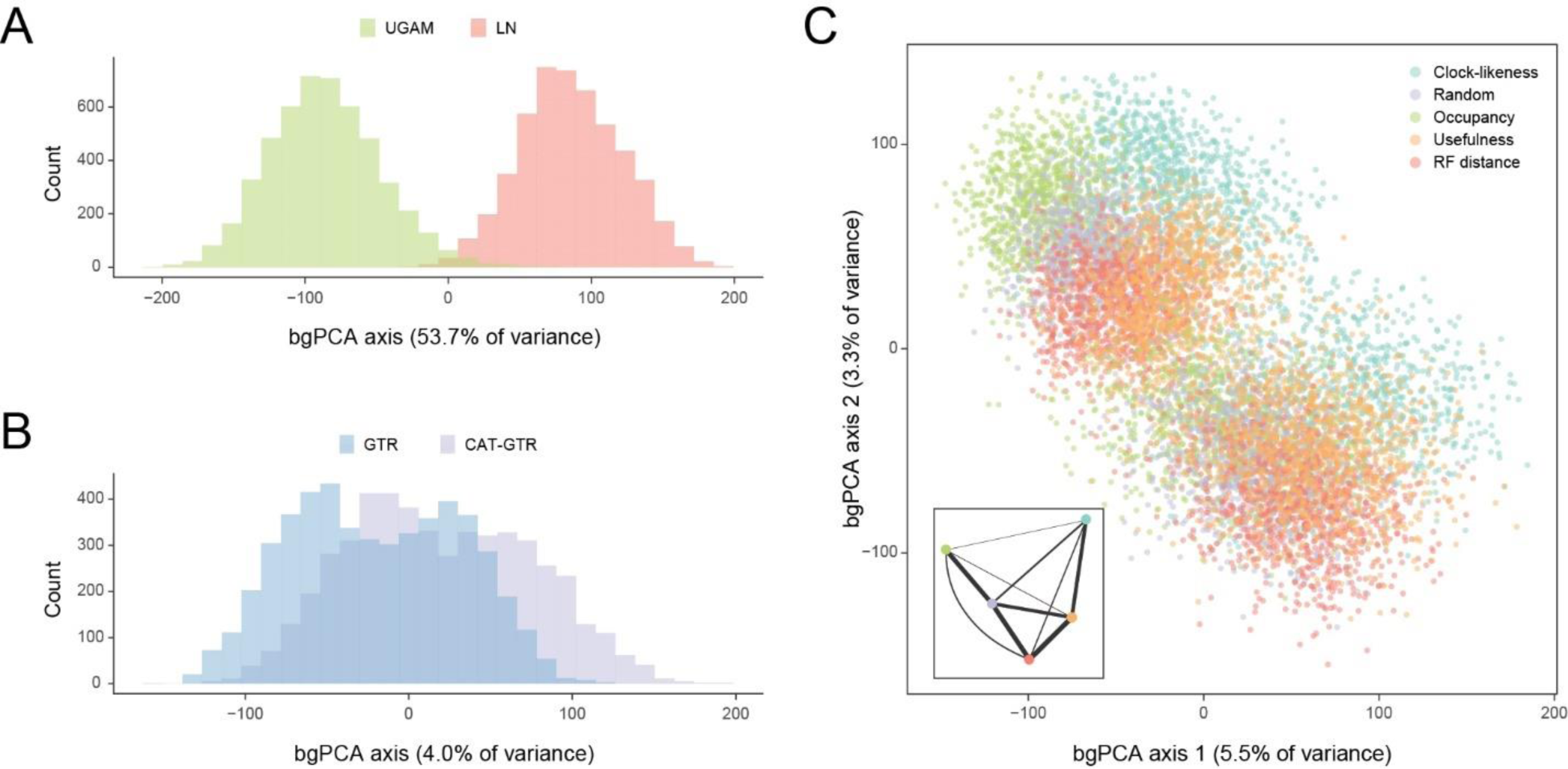
Sensitivity of divergence time estimation to methodological decisions. Between-group principal component analysis (bgPCA) was used to retrieve axes that separate chronograms based on the clock model (**A**), model of molecular evolution (**B**), and gene sampling strategy (**C**) employed. In the latter case, only the first two out of four bgPCA dimensions are shown. The inset shows the centroid for each loci sampling strategy, and the width of the lines connecting them are scaled to the inverse of the Euclidean distances that separates them (as a visual summary of overall similarity). The proportions of total variance explained are shown on the axis labels. The impact of the clock model is such that a bimodal distribution of chronograms can be seen even when bgPCA are built to discriminate based on other factors (as in **C**).

Regarding gene choice, the ages most different to those obtained under random loci selection were found when using the most clock-like genes (Fig. 3C).

### **(c)** Echinoid (and echinoderm) divergence times

Even when the age of crown Echinodermata was constrained to postdate the appearance of stereom (the characteristic skeletal microstructure of echinoderms) in the Early Cambrian [42, 43], only analyses using the most clock-like loci recovered ages concordant with this (i.e., median ages younger than the calibration enforced; Fig. S4). Instead, most consensus trees favored markedly older ages for the clade, in some cases even predating the origin of the Ediacaran biota [44] (Fig. S6). Despite the relative sensitivity of many of the earliest nodes to methodological choices (Fig. S2), the split between Crinoidea and all other echinoderms (Eleutherozoa) is always inferred to have predated the end of the Cambrian (youngest median age = 492.2 Ma), and the divergence among the other major lineages (classes) of extant echinoderms are constrained to have happened between the Late Cambrian and Middle Ordovician (Fig. S6). Our results also recover an early origin of crown group Holothuroidea (sea cucumbers; range of median ages = 351.6–383.2 Ma), well before the crown groups of other extant echinoderm classes. These dates markedly postdate the first records of holothuroid calcareous rings in the fossil record [45, 46], and imply that this trait does not define the holothuroid crown group but instead evolved from an echinoid-like jaw-apparatus along its stem [47]. The other noteworthy disagreement between our results and those of previous studies [48] involves dating crown group Crinoidea to times that precede the P-T mass extinction (range of median ages = 268.0–327.7 Ma, although highest posterior density intervals are always wide and include Triassic ages).

Across all of the analyses performed, the echinoid crown group is found to have originated somewhere between the Pennsylvanian and Cisuralian, with 42% posterior probability falling within the late Carboniferous and 58% within the early Permian (Fig. 4). An origin of the clade postdating the P-T mass extinction is never recovered, even when such ages were allowed under the joint prior (Fig. S7).

**Figure 4:**
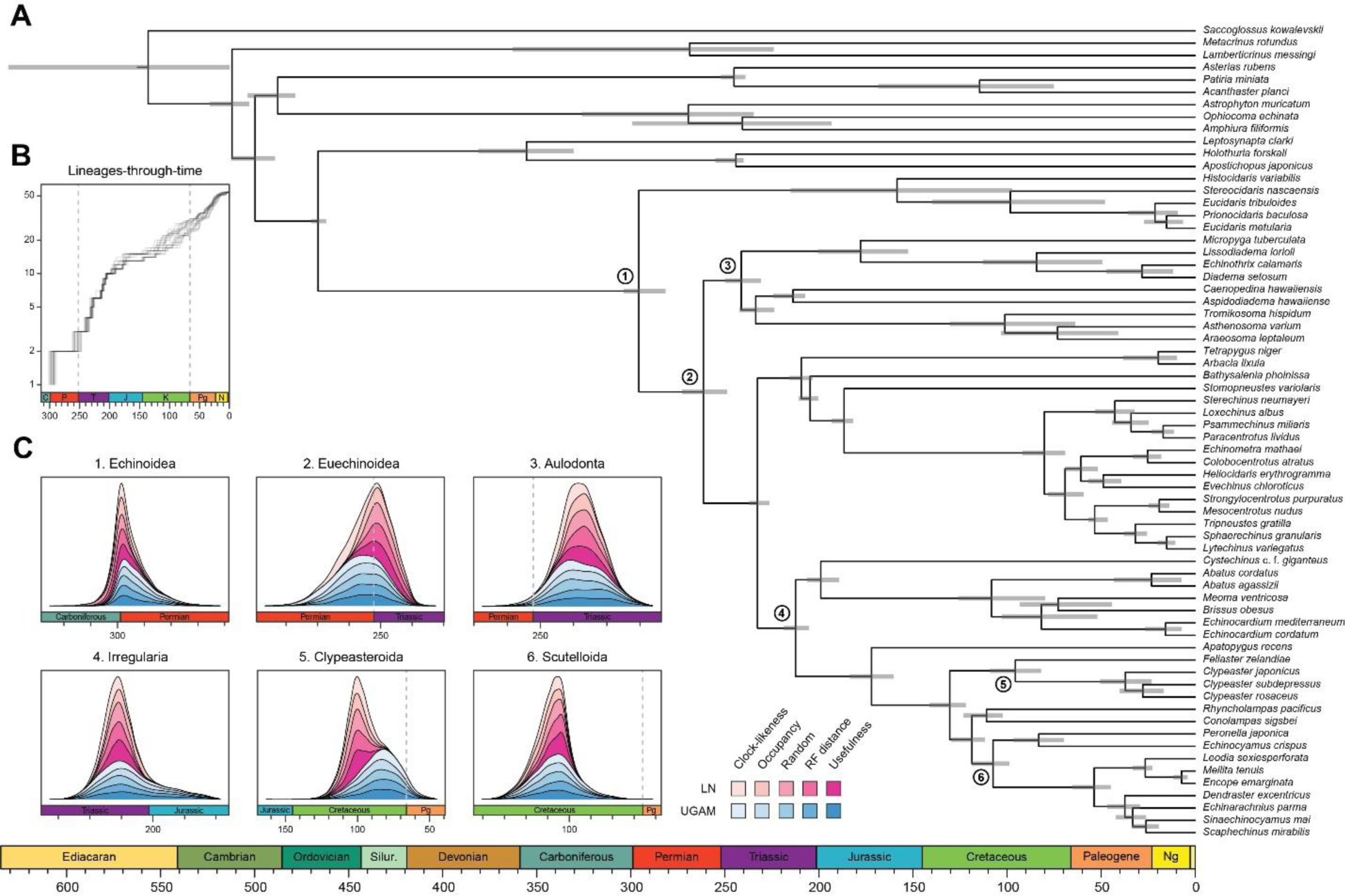
Divergence times among major clades of Echinoidea and other echinoderms. **A.** Consensus chronogram of the two runs using clock-like genes under a CAT+GTR+G model of molecular evolution and a autocorrelated log-normal (LN) clock prior. Node ages correspond to median values, and bars show the 95% highest posterior density intervals. **B.** Lineage-through-time plot, showing the rapid divergence of higher-level clades following the P-T mass extinction (shown with dashed lines, along with the K-Pg boundary). Each line corresponds to an individual consensus topology from among the time- calibrated runs performed. **C.** Posterior distributions of the ages of selected nodes (identified in **A**). The effects of models of molecular evolution are not shown, as they represent the least important factor (see Fig. 3); distributions using site-homogeneous and heterogeneous models are combined for every combination of targeted loci and clock prior.

While the posterior distribution of ages for Euechinoidea spans both sides of the P-T boundary, the remaining earliest splits within the echinoid tree are constrained to have occurred during the Triassic, including the origins of Aulodonta, Carinacea, Echinacea, and Irregularia (Figs. 4 and S6). Many echinoid orders are also inferred to have diverged from their respective sister clades during this period, including aspidodiadematoids, pedinoids, echinothurioids, arbacioids, and salenioids. LTT plots confirm that lineage diversification proceeded rapidly throughout the Triassic (Fig. 4B). Despite the topological reorganization of Neognathostomata, the clade is dated to a relatively narrow time interval in the Late to Middle Jurassic (range of median ages = 169.48-179.96 Ma), in agreement with recent estimates [3]. Within this clade, the origins of both scutelloids and clypeasteroids confidently predate the K-Pg mass extinction (posterior probability of origination before the boundary = 0.9995 and 0.9621, respectively), despite younger ages being allowed by the joint prior (Fig. S7).

## Discussion

### **(a)** The echinoid Tree of Life

In agreement with previous phylogenomic studies [3, 4], echinoid diversity can be subdivided into five major clades (Fig. 2A). Cidaroids form the sister group to all other crown group echinoids (Euechinoidea). Some aspects of the relationships among sampled cidaroids are consistent with previous molecular [49] and morphological studies [19], including an initial split between *Histocidaris* and the remaining taxa, representing the two main branches of extant cidaroids [5, 35]. Others, such as the nested position of *Prionocidaris baculosa* within the genus *Eucidaris,* not only implies paraphyly of this genus but also suggests the need for a taxonomic reorganization of the family Cidaridae. Within euechinoids, the monophyly of Aulodonta is supported for the first time with sampling of all of its major groups. The subdivision of these into a clade that includes diadematoids plus micropygoids (which we propose should retain the name Diadematacea), sister to a clade including echinothurioids and pedinoids (Echinothuriacea *sensu* [4]) is strongly reminiscent of some early classifications (e.g., [50]).

Our expanded phylogenomic sampling also confirms an aulodont affinity for aspidodiadematoids [3, 35] and places them within Echinothuriacea as the sister group to Pedinoida.

In agreement with previous phylogenomic studies [3, 4], echinoid diversity can be subdivided into five major clades (Fig. 2A). Cidaroids form the sister group to all other crown group echinoids (Euechinoidea). Some aspects of the relationships among sampled cidaroids are consistent with previous molecular [47] and morphological studies [19], including an initial split between *Histocidaris* and the remaining taxa, representing the two main branches of extant cidaroids [5, 35]. Others, such as the nested position of *Prionocidaris baculosa* within the genus *Eucidaris,* not only implies paraphyly of this genus but also suggests the need for a taxonomic reorganization of the family Cidaridae. Within euechinoids, the monophyly of Aulodonta is supported for the first time with sampling of all of its major groups. The subdivision of these into a clade that includes diadematoids plus micropygoids (which we propose should retain the name Diadematacea), sister to a clade including echinothurioids and pedinoids (Echinothuriacea *sensu* [4]) is strongly reminiscent of some early classifications (e.g., [48]).

Our expanded phylogenomic sampling also confirms an aulodont affinity for aspidodiadematoids [3, 35] and places them within Echinothuriacea as the sister group to Pedinoida.

The remaining diversity of echinoids, which forms the clade Carinacea (Fig. 2), is subdivided into Irregularia and their sister clade among regulars, for which we amend the name Echinacea to include Salenioida. Given the striking morphological gap separating regular and irregular echinoids, the origin of Irregularia has been shrouded in mystery [19, 23, 50]. Our complete sampling of major regular lineages determines Echinacea *sensu stricto* to be the sister clade to irregular echinoids. A monophyletic Echinacea was also supported in a recent total-evidence analysis [3], but the incomplete molecular sampling of that study resulted in a slightly different topology that placed salenioids as the sister group to the remaining lineages. However, an overall lack of morphological synapomorphies uniting these clades had previously been acknowledged [19]. While the relationships within Echinacea proved to be difficult to resolve even with thousands of loci, multiple lines of evidence lead us to prefer a topology in which salenioids form a clade with camarodonts + stomopneustoids, with arbacioids sister to all of these (Fig. 2).

As has been already established [3, 4, 14, 19, 30], the lineages of irregular echinoids here sampled are subdivided into Atelostomata (heart urchins and allies) and Neognathostomata (sand dollars, sea biscuits, and “cassiduloids”). Despite the former being the most diverse of the five main clades of echinoids (Fig. 2), its representation in phylogenomic studies remains low, and its internal phylogeny poorly constrained [35]. On the contrary, recent molecular studies have greatly improved our understanding of the relationships among neognathosomates [3, 4, 36], revealing an evolutionary history that dramatically departs from previous conceptions. Even when scutelloids and clypeasteroids were never recovered as reciprocal sister lineages by molecular phylogenies (e.g., [13, 14, 30]), this result was not fully accepted until phylogenomic data confidently placed echinolampadids as the sister lineage to sand dollars [4]. At the same time, this result rendered the position of the remaining “cassiduloids”, a taxonomic wastebasket with an already complicated history of classification [19, 27, 28, 51], entirely uncertain. An attempt to constrain the position of these using a total-evidence approach [3] subdivided the “cassiduloids“ into three unrelated clades: Nucleolitoida, composed of extinct lineages and placed outside the node defined by Scutelloida + Clypeasteroida, and two other clades nested within it (see Fig. 1). Extant “cassiduloids” were recovered as members of one of the latter clades, representing the monophyletic sister group to sand dollars. Here, we show that *Apatopygus recens* does not belong within this clade but is instead the sister group to all other extant neognathostomates. Given this phylogenetic position, as well as the morphological similarities between *Apatopygus* and the entirely extinct nucleolitids [19, 28, 51–53], it is likely that the three extant species of apatopygids represent the last surviving remnants of Nucleolitoida, a clade of otherwise predominantly Mesozoic neognathostomates [3]. Because of the renewed importance in recognizing this topology, we propose the name Luminacea for the clade uniting all extant neognathostomates with the exclusion of *Apatopygus* (Fig. 2A). This nomenclature refers to the dynamic evolutionary history of the Aristotle’s lantern (i.e., the echinoid jaw-apparatus) within the clade (present in the adults of both clypeasteroids and scutelloids, but found only in the juveniles of Cassiduloida *sensu stricto*), the inclusion of the so-called lamp urchins (echinolampadids) within the clade, and the illumination provided by this hitherto unexpected topology. The previous misplacement of *Apatopygus* within this clade ([3]; see Fig. 1G) is likely a consequence of tip-dating preferring more stratigraphically congruent topologies [54], an effect that can incorrectly resolve taxa on long terminal branches [55]. Given the generally useful phylogenetic signal of stratigraphic information [56], this inaccuracy further highlights the unusual evolutionary history of living apatopygids.

### **(b)** Chronospaces: a statistical exploration of time-calibration strategies

Calibrating phylogenies to absolute time is crucial to understanding evolutionary history, as the resulting chronograms provide a major avenue for testing hypotheses of diversification, character evolution and other macroevolutionary processes. However, the accuracy and precision of the inferred divergence times hinge upon many methodological choices that are often difficult or time-consuming to justify (e.g., calibration strategies, prior distributions on node ages, clock models, etc.) [57–62].

Understanding the impact of each of these sources of uncertainty can be difficult. The sensitivity of inferred ages is commonly explored by running analyses under different settings and summarizing the results in tables or by stacking chronograms in order to visualize the relative position of nodes (see for example [58, 59]). This approach discards a lot of information contained in Bayesian posterior distributions of time-calibrated topologies. For example, patterns of correlation among branch lengths (i.e., ways in which changes in a given branch length are accommodated by other regions of the tree) are lost, and alternative ways in which node ages are attained (e.g., which of all descendant branches expand as a given node becomes older) are not usually explored, even though they can potentially represent very different evolutionary scenarios.

Here we analyze the sensitivity of node ages to alternative criteria to sample loci from phylogenomic datasets, as well as different ways of modelling the patterns of molecular evolution across sites and the rates of evolution among lineages. To do so, we introduce an approach to visualize the distribution of chronograms in a multidimensional space of node ages, a chronospace, and measure the overall effect of these decisions on inferred dates using multivariate statistical methods. Our results reveal that divergence times obtained under site-homogeneous and site-heterogeneous models (such as CAT+GTR+G) are broadly comparable. This happens despite the latter estimating higher levels of sequence divergence and stretching branches in a non-isometric manner (Fig. S1). Site-heterogeneous models have become common for the analysis of phylogenomic datasets spanning deep divergences, as they exhibit reduced sensitivity to saturation and are thus more robust to long-branch attraction artifacts [63–65]. These models are also central to the debate regarding contentious nodes in the tree of life, including the position of ctenophores [66, 67] and the monophyly of arachnids [68, 69]. While the use of CAT models for time calibration has become equally popular, the degree to which they impact estimates of node ages has received less scrutiny. The lack of a meaningful effect uncovered here, coupled with their high computational burden [70], questions their usefulness for time-scaling phylogenies. A similar result was recently found when comparing site-homogeneous models with different numbers of parameters [71], suggesting that relaxed clocks adjust branch rates in a manner that buffers the effects introduced by using more complex (and computationally-intensive) models of sequence evolution.

Similarly, the choice between different loci has a small effect on inferred ages, with little evidence of a systematic difference between the divergence times supported by randomly-chosen loci and those found using targeted sampling criteria, such as selecting genes for their phylogenetic signal, usefulness, occupancy, or clock-likeness. A meaningful effect was restricted to a few ancient nodes (e.g., Echinodermata), for which clock-like genes suggested younger ages that are more consistent with fossil evidence. This result provides direct empirical evidence that validates the selection of clock-like genes for inferring deep histories of diversification, an approach so far supported mostly by simulations [62, 72]. At the same time, the choice of loci seems to be less relevant for younger nodes, at least when employing large numbers of characters as done here. Finally, the choice between alternative clock models induced differences in ages that were five to ten times stronger than those of other factors, emphasizing the importance of either validating their choice (e.g., [73]) or—as done here—focusing on results that are robust to them.

### **(c)** Echinoid origins and diversification

The origin and early diversification of crown group Echinoidea have always been considered to have been determined (or at least strongly affected) by the P-T mass extinction [11, 12, 16, 20].

However, estimating the number of crown group members surviving the most severe biodiversity crisis in the Phanerozoic [74] has been hampered by both paleontological and phylogenetic uncertainties [3, 10, 13–15, 18]. Our results establish with confidence that multiple crown group lineages survived and crossed this boundary, finding for the first time a null posterior probability of the clade originating after the extinction event. While the survival of three crown group lineages is slightly favored (Fig. S8), discerning between alternative scenarios is still precluded by uncertainties in dating these early divergences. Echinoid diversification during the Triassic was relatively fast (Fig. 4B) and involved rapid divergences among its major clades. Even many lineages presently classified at the ordinal level trace their origins to this initial pulse of diversification following the P-T mass extinction.

The late Paleozoic and Triassic origins inferred for the crown group and many euechinoid orders prompts a re-evaluation of fossils from this interval of time. Incompletely known fossil taxa such as the Pennsylvanian *?Eotiaris meurevillensis*, with an overall morphology akin to that of crown group echinoids, has a stratigraphic range consistent with our inferred date for the origin of the echinoid crown group [20]. Additionally, the Triassic fossil record of echinoids has been considered to be dominated by stem group cidaroids, with the first euechinoids not known until the Late Triassic [17, 75]. However, the Triassic origins of many euechinoid lineages supported by our analyses necessitates that potential euechinoid affinities should be re-considered for this diversity of Triassic fossils. This is especially the case for the serpianotiarids and triadocidarids, abundant Triassic families variously interpreted as cidaroids, euechinoids, or even stem echinoids [3, 17, 75, 76]. A reinterpretation of any of these as euechinoids would suggest that the long-implied gap in the euechinoid record [16, 18] is caused by our inability to correctly place these key fossils, as opposed to an incompleteness of the fossil record itself.

While our phylogenomic approach is the first to resolve the position of all major cassiduloid lineages, the inferred ages for many nodes within Neognathostomata remain in strong disagreement with the fossil record. No Mesozoic fossil can be unambiguously assigned to either sand dollars or sea biscuits, a surprising situation given the good fossilization potential and highly distinctive morphology of these clades [19, 21, 25, 77]. While molecular support for a sister group relationship between scutelloids and echinolampadids already implied this clade (Echinolampadacea) must have split from clypeasteroids by the Late Cretaceous [3, 4, 14, 19], this still left open the possibility that the crown groups of sand dollars and sea biscuits radiated in the Cenozoic. Under this scenario, the Mesozoic history of these groups could have been comprised of forms lacking their distinctive morphological features, complicating their correct identification. This hypothesis is here rejected, with the data unambiguously supporting the radiation of both crown groups preceding the K-Pg mass extinction (Fig. 4C). While it remains possible that these results are incorrect even after such a thorough exploration of the time-calibration toolkit (see for example [62, 78]), these findings call for a critical reassessment of the Cretaceous fossil record, and a better understanding of the timing and pattern of morphological evolution among fossil and extant neognathostomates. For example, isolated teeth with an overall resemblance to those of modern sand dollars and sea biscuits have been found in Lower Cretaceous deposits [79], raising the possibility that other overlooked and disarticulated remains might close the gap between rocks and clocks.

### **(d)** Conclusions

Although echinoid phylogenetics has long been studied using morphological data, the position of several major lineages (e.g., aspidodiadematoids, micropygoids, salenioids, apatopygids) remained to be confirmed with the use of phylogenomic approaches. Our work not only greatly expands the available genomic resources for the clade, but finds novel resolutions for some of these lineages, improving our understanding of their evolutionary history. The most salient aspect of our topology is the splitting of the extant “cassiduloids” into two distantly related clades, one of which is composed exclusively of apatopygids. This result is crucial to constrain the ancestral traits shared by the main lineages of neognathostomates, helping unravel the evolutionary processes that gave rise to the unique morphology of the sand dollars and sea biscuits [3, 24, 25].

Although divergence time estimation is known to be sensitive to many methodological decisions, systematically quantifying the relative impact of these on inferred ages has rarely been done. Here we propose an approach based on chronospaces (implemented in R [80] and available at https://github.com/mongiardino/chronospace) that can help visualize key effects and determine the sensitivity of node dates to different assumptions. Our results shed new light on the early evolutionary history of crown group echinoids and its relationship with the P-T mass extinction event, a point in time where the fossil record provides ambiguous answers. They also establish with confidence a Cretaceous origin for the sand dollars and sea biscuits, preceding their first appearance in the fossil record by at least 40 to 50 Ma, respectively (and potentially up to 65 Ma). These clades, therefore, join several well-established cases of discrepancies between the fossil record and molecular clocks, such as those underlying the origins of placental mammals [81] and flowering plants [82].

## Methods

### **(a)** Sampling, bioinformatics, and matrix construction

This study builds upon previous phylogenomic matrices [3, 4], the last of which was augmented through the addition of eight published datasets (mostly expanding outgroup sampling), as well as 17 novel echinoid datasets (15 transcriptomes and two draft genomes). For all novel datasets, tissue sampling, DNA/RNA extraction, library preparation, and sequencing varied by specimen, and are detailed in the Supplementary Information. Raw reads have been deposited in NCBI under Bioproject accession numbers XXX and XXX. Final taxon sampling included 12 outgroups and 50 echinoids (accession numbers and sampling details can be found in Table S1).

Raw reads for all transcriptomic datasets were trimmed or excluded using quality scores with Trimmomatic v. 0.36 [83] under default parameters. Further sanitation steps were performed using the Agalma 2.0 phylogenomic workflow [84], and datasets were assembled *de novo* with Trinity 2.5.1 [85]. For genomic shotgun sequences, adapters were removed with BBDuk (https://sourceforge.net/projects/bbmap/), and UrQt v. 1.0.18 [86] was used to filter short reads (size < 50) and trim low-quality ends (score < 28). Datasets were then assembled using MEGAHIT v. 1.1.2 [87].

Draft genomes were masked using RepeatMasker v. 4.1.0 [88, 89], before obtaining gene predictions with AUGUSTUS 3.2.3 [90]. A custom set of universal single-copy orthologs (USCOs) obtained from the latest *Strongylocentrotus purpuratus* genome assembly (Spur v. 5.0) was employed as the training dataset. Settings and further details of these analyses can be found in the Supplementary Information.

Multiplexed transcriptomes were sanitized from cross-contaminants using CroCo v. 1.1 [91], and likely non-metazoan contaminants were removed using alien_index v. 3.0 [92] (removing sequences with AI values > 45). Datasets were imported back into Agalma, which automated orthology inference (as described in [84, 93]), gene alignment with MACSE [94], and trimming with GBLOCKS [95]. The amino acid supermatrix was reduced using a 70% occupancy threshold, producing a final dataset of 1,346 loci (327,695 sites). As a final sanitation step, gene trees were obtained using ParGenes v. 1.0.1 [96], which performed model selection (minimizing the Bayesian Information Criterion) and inference using 100 bootstrap (BS) replicates. Trees were then used to remove outlier sequences with TreeShrink v. 1.3.1 [97]. We specified a reduced tolerance for false positives and limited removal to at most three terminals which had to increase tree diameter by at least 25% (-q 0.01 -k 3 -b 25). Statistics for the supermatrix and all assemblies can be found in Table S2.

### **(b)** Phylogenetic inference

Inference was performed under multiple probabilistic and coalescent-aware methods, known to differ in their susceptibility to model violations. Coalescent-based inference was performed using the summary method ASTRAL-III [98], estimating support as local posterior probabilities [99]. Among concatenation approaches, we used Bayesian inference under an unpartitioned GTR+G model in ExaBayes 1.5 [100]. Two chains were run for 2.5 million generations, samples were drawn every one hundred, and the initial 25% was discarded as burn-in. We also explored maximum likelihood inference with partitioned and unpartitioned models. For the former, the fast-relaxed clustering algorithm was used to find the best-fitting model among the top 10% using IQ-TREE 1.6.12 [101, 102] (-m MFP+MERGE -rclusterf 10 -rcluster-max 3000), and support was evaluated with 1,000 ultrafast BS replicates [103]. For the latter, we used the LG4X+R model in RAxML-NG v. 0.5.1 [104] and evaluated support with 200 replicates of BS. Finally, we also implemented the site-heterogeneous LG+C60+F+G mixture model using the posterior mean site frequency (PMSF) approach to provide a fast approximation of the full profile mixture model [105], allowing the use of 100 BS replicates to estimate support. Given some degree of topological conflict between the results of the other methods (see below), multiple guide trees were used to estimate site frequency profiles, but the resulting phylogenies were identical.

Given conflicts between methods in the resolution of one particular node (involving the relationships among Arbacioida, Salenioida, and the clade of Stomopneustoida + Camarodonta), all methods were repeated after reducing the matrix to 500 and 100 loci selected for their phylogenetic usefulness using the approach described in [3, 37] and implemented in the *genesortR* script (https://github.com/mongiardino/genesortR). This approach relies on seven gene properties routinely used for phylogenomic subsampling, including multiple proxies for phylogenetic signal—such as the average BS and Robinson-Foulds (RF) similarity to a target topology—as well as several potential sources of systematic bias (e.g., rate and compositional heterogeneities). Outgroups were removed before calculating these metrics. RF similarity was measured to a species tree that had the conflicting relationship collapsed so as not to bias gene selection in favor of any resolution. A principal component analysis (PCA) of this dataset resulted in a dimension (PC 2, 17.6% of variance) along which phylogenetic signal increased while sources of bias decreased (Fig. S9), and which was used for loci selection. For the smallest dataset, we also performed inference under the site-heterogeneous CAT+GTR+G model using PhyloBayes-MPI [106]. Three runs were continued for > 10,000 generations, sampling every two generations and discarding the initial 25%. Convergence was confirmed given a maximum bipartition discrepancy of 0.067 and effective sample sizes for all parameters > 150.

Two other approaches were implemented in order to assist in resolving the contentious node.

First, we implemented a likelihood-mapping analysis [107] in IQ-TREE to visualize the phylogenetic signal for alternative resolutions of the quartet involving these three lineages (Arbacioida, Salenioida, and Stomopneustoida + Camarodonta) and their sister clade (Irregularia; other taxa were excluded). Second, we estimated the log-likelihood scores of each site in RAxML (using best-fitting models) for the two most strongly supported resolutions found through likelihood mapping. These were used to calculate gene-wise differences in scores, or δ values [108]. In order to search for discernable trends in the signal for alternative topologies, genes were ordered based on their phylogenetic usefulness (see above) and the mean per-locus δ values of datasets composed of multiples of 20 loci (i.e., the most useful 20, 40, etc.) were calculated.

### **(c)** Time calibration

Node dating was performed using relaxed molecular clocks in PhyloBayes v4.1 using a fixed topology and a novel set of 22 fossil calibrations corresponding to nodes from our newly inferred phylogeny (listed in the Supplementary Information). Depending on the node, we enforced both minimum and maximum bounds, or either one of these. A birth-death prior was used for divergence times, which allowed for the implementation of soft bounds [109], leaving 5% prior probability of divergences falling outside of the specified interval. We explored the sensitivity of divergence time estimates to gene selection, model of molecular evolution, and clock prior. One hundred loci were sampled from the full supermatrix according to four targeted sampling schemes: usefulness (calculated as explained above, except incorporating all echinoderm terminals), phylogenetic signal (i.e., smallest RF distance to species tree), clock-likeness (i.e., smallest variance of root-to-tip distances), and level of occupancy. For clock-likeness, we only considered loci that lay within one standard deviation of the mean rate (estimated by dividing total tree length by the number of terminals [110]), as this method is otherwise prone to selecting largely uninformative loci (Fig. S10; [37]). A fifth sample of randomly chosen loci was also evaluated. These five datasets were run under two models of molecular evolution, the site-homogeneous GTR+G and the site-heterogeneous CAT+GTR+G, and both uncorrelated gamma (UGAM) and autocorrelated log-normal (LN) clocks were implemented. The combination of these settings (loci sampled, model of evolution, and clock prior) resulted in 20 analyses. For each, two runs were continued for 20,000 generations, after which the initial 25% was discarded and the chains thinned to every two generations (see log-likelihood trace plots in the electronic supplementary material, Fig. S11). To explore the sensitivity of divergence times to these methodological decisions, 500 random chronograms were sampled from each analysis (250 from each run), and their node dates were subjected to multivariate analyses of variance (MANOVA) and between-group PCA (bgPCA) using package *Morpho* [111] in the R statistical environment [80]. bgPCA involves the use of PCA on the covariance matrix of group means (e.g., the mean ages of all nodes obtained under UGAM or LN clocks), followed by the projection of original samples onto the obtained bgPC axes. The result is a multidimensional representation of divergence times—a chronospace—rotated so as to capture the distinctiveness of observations obtained under different settings. Separate bgPCAs were performed for loci sampling strategy, clock prior and model of molecular evolution, and the proportion of total variance explained by bgPC axes was interpreted as an estimate of the relative impact of these choices on inferred times of divergence. Finally, lineage-through-time plots were generated using *ape* [112].

## Author contributions

NMK, RM, and GWR conceived and designed the study. NMK, ASH, MFM, AFA, SEC, FA, OB, and AK performed extractions, prepared libraries, and sequenced transcriptomic/genomic datasets. NMK and AK generated *de novo* assemblies. JRT and RM designed fossil calibrations. NMK run all phylogenetic analyses. NMK and JRT wrote the manuscript with input from all other authors. All authors gave final approval for publication and agree to be held accountable for the work performed therein.

## Data, code and materials

Raw transcriptomic reads have been deposited in NCBI under Bioproject accession number XXX. Assemblies are available from the authors upon request. R code to plot chronospaces and explore the effects of methodological decisions on divergence time estimation can be found at https://github.com/mongiardino/chronospace.

## Acknowledgments

We are grateful to the crew of R/Vs Antéa, Atlantis, Falkor and Mirai, HOV Alvin, and the crew and PIs of BIOMAGLO deep-sea cruises (Laure Corbari, Karine Olu-Le Roy, Sarah Samadi) for assistance in specimen collection. We would also like to thank Libby Liggins, Wilma Blom, Owen Anderson, Francisco Solís-Marín, Carlos A. Conejeros-Vargas, and Jih-Pai Lin for the collection and shipment of specimens.

We gratefully acknowledge assistance from Thomas Saucède (Univ. Burgundy), Marc Eléaume (MNHN), and Charlotte Seid (Scripps) in accessing, cataloging and processing museum specimens. Tim Ravasi provided resources and collection facilities to GWR via King Abdullah University of Science and Technology (KAUST). Institutional support was provided by the Central Research Laboratories and the Department of Geology and Palaeontology at the Natural History Museum Vienna, Austria. This work was supported by a Yale Institute for Biospheric Studies Doctoral Dissertation Improvement Grant and a Society of Systematic Biologists Graduate Student Research Award to NMK. AK received funding from an Austrian Science Fund project (FWF, project number P 29508-B25), FA from Agencia Nacional de Investigación (ANID/PAI Inserción en la Academia, project number PAI79170033), and GWR and RM from NSF grant DEB-2036186. JRT was supported by a Royal Society Newton International Fellowship and a Leverhulme Trust Early Career Fellowship. NMK was supported by a Yale University fellowship.

## Supplementary Information

### Sequencing and further bioinformatic details

Apatopygus recens, Aspidodiadema hawaiiense, Fellaster zelandiae, Histocidaris variabilis, Rhyncholampas pacificus, Sinaechinocyamus mai, Stereocidaris neumayeri, Tromikosoma hispidum. Tissue subsamples were finely chopped with a scalpel and preserved in RNAlater (Ambion) buffer solution for 1 day at 4°C to allow the RNAlater to effectively penetrate the tissues, followed by long-term storage at -80°C until RNA extraction. Total RNA was extracted from 1.5 mm Triple-Pure High Impact Zirconium beads (Benchmark Scientific) in Trizol (Ambion), using Direct-zol RNA Miniprep Kit (Zymo Research) with in-column DNase I incubation to remove genomic DNA. Prior to mRNA capture, total RNA concentration was estimated using Qubit RNA HS Assay Kit (Invitrogen; range = 8.33-96 ng/μL), and quality was assessed using either High Sensitivity RNA ScreenTape or RNA ScreenTape with an Agilent 4200 TapeStation. Mature mRNA was isolated from total RNA and libraries were prepared using KAPA mRNA HyperPrep Kit (KAPA Biosystems) following the manufacturer’s instructions (including sample customization based on total RNA quantity and quality values), targeting an insert size circa 500 base pairs (bp), and using custom 10-nucleotide Illumina TruSeq style adapters [113]. Post-amplification, DNA concentration was estimated using Qubit dsDNA BR Assay Kit (Invitrogen; range = 6.06-13.4 ng/μL). Concentration, quality, and molecular weight distribution of libraries (range = 547-978 bp) was also assessed using Genomic DNA ScreenTape with an Agilent 4200 TapeStation. Ten libraries (including two annelids not employed here) were sequenced on one lane of a multiplexed run using NovaSeq S4 platform with 100 bp paired-end reads at the IGM Genomics Center (University of California San Diego).

The assembled transcriptomes were sanitized from cross-contaminant reads product of multiplexed sequencing using CroCo v. 1.1 [91]. These eight datasets, as well as two annelid transcriptomes not employed in this study but sequenced in the same lane, were incorporated. Transcripts considered over or under-expressed across samples (as defined by default parameters), were kept. This resulted in an average removal of 1.26% of assembled transcripts (range: 0.4% for *Aspidodiadema*to 3.06% for *Histocidaris*).

*Clypeaster japonicus*, *Encope emarginata*, *Leodia sexiesperforata*, *Peronella japonica*. Eggs were collected from a single female specimen of each species and immediately placed into RNA later for preservation. The samples were left in RNAlater at 4°C for at least 1 day and then transferred to a -80°C freezer until RNA extraction. RNA extraction was performed using Trizol and was then treated with Ambion’s Turbo DNA-free kit. RNA was quantified using both Nanodrop and an Agilent 2100 Bioanalyzer. Only RNA samples with an RNA integrity number (RIN) of > 7 were used. RNA-Seq libraries were prepared using the NEB Ultra Directional kit using the standard protocol and multiplexed with 6bp NEB multiplex primers. Paired-end 100 libraries were generated from the resultant libraries at the UC Berkeley Genome Center using an Illumina HighSeq 2500.

*Bathysalenia phoinissa*, *Micropyga tuberculata*: The material was collected by the deep-sea cruise BIOMAGLO [114] conducted in 2017 jointly by the French National Museum of Natural History (MNHN) as part of the Tropical Deep-Sea Benthos program, the French Research Institute for

Exploitation of the Sea (IFREMER), the “Terres Australes et Antarctiques Françaises” (TAAF), the Departmental Council of Mayotte and the French Development Agency (AFD), with the financial support of the European Union (Xe FED). Specimens were sorted into taxonomic bins at class level and collectively fixed in 70% ethanol at room temperature. Following taxonomic identification, tissue samples were obtained using sterilized forceps and disposable scalpel blades. Tissue subsamples were collected in high-purity 96% ethanol and stored at −40°C until DNA extraction. Total genomic DNA was extracted using the DNeasy Blood and Tissue Kit (QIAGEN) following the manufacturer’s instructions, complemented by the addition of 4 µl RNAse A (QIAGEN, 100 mg/ml) for RNA digestion. Elution buffer volume was reduced to 60 µl and the elution step repeated twice per sample in order to maximize DNA concentration and yield. After collection of the first elution, spin-column membranes were rinsed again twice with fresh elution buffer (40 µl) to further increase yield. Elutions were collected separately, with the first one used for sequencing and the second for quality control. For the latter, a portion of the mitochondrial cytochrome c oxidase subunit I (COI) was amplified using PCR. Amplifications were conducted with TopTaq DNA Polymerase (QIAGEN) using 1 μl of extracted genomic DNA (approx. 10–15 ng). Details on primers and protocols can be found in [115]. PCR products were visualized on a 1% agarose gel, purified using ExoSAP-IT (Affymetrix), and sequenced at Microsynth GmbH (Vienna, Austria) with the same primers. Sequences are deposited in GenBank under the accession numbers XXX and XXX.

Prior to library preparation, total genomic DNA concentration was estimated using Qubit DNA BR Assay Kit (Invitrogen) in order to determine library preparation strategy. This step was carried out by Macrogen (Korea), using an Illumina TruSeq DNA PCR-free kit (for *Micropyga*) and an Illumina TruSeq Nano DNA kit (for *Bathysalenia*). Library preparation followed the manufacturer’s instructions, targeting an insert size of 350 base pairs (bp). The libraries were sequenced by Macrogen (Korea) on a shared lane of a multiplexed run using an Illumina HiSeq X instrument with 150 bp paired-end reads. Sequencing depth was 134.1 and 161.3 million reads for *Bathysalenia* and *Micropyga*, respectively. Pre-processing of Illumina shotgun data was carried out by removal of adapters using BBDuk (https://sourceforge.net/projects/bbmap/) with settings: `minlen=50 ktrim=r k=23 mink=11 tpe tbo`); followed by quality trimming of reads using UrQt v. 1.0.18 [86], discarding regions with phred quality score < 28 and sequences of size < 50. Assembly of the data was carried out with MEGAHIT v. 1.1.2 [87], in an iterative approach followed by assessment of assembly quality using BUSCO v. 3.0.2. scores [116] using the metazoan dataset ODB v. 9 [117]. Final BUSCO scores of the draft assemblies were 28.4% (S:27.8%, D:0.6%, F:48.2%, M:23.4%, n:978) for *Bathysalenia* and 17.2% (S:16.8%, D:0.4%, F:59.0%, M:23.8%, n:978) for *Micropyga*. As recommended by Hoff and Stanke [89], draft genomes were masked with RepeatMasker v. 4.1.0 [88] prior to gene prediction, using settings: `-norna -xsmall`; resulting in 3.82 % and 2.9% masked bases for *Bathysalenia* and *Micropyga*, respectively. Gene prediction was carried out using AUGUSTUS v. 3.2.3 [90] using settings: `--strand=both -- singlestrand=false --genemodel=partial --codingseq=on --sample=0 -- alternatives-from-sampling=false --exonnames=on --softmasking=on`. This step relied on a training dataset composed of universal single-copy orthologs (USCOs) derived from the most recent *Strongylocentrotus purpuratus* genome (Spur v.5).

*Echinothrix calamaris*, *Tripneustes gratilla*. Tissue samples were taken from live sea urchins housed in laboratory aquaria and total RNA was immediately extracted from 150 mg of tissue using a PureLink™ RNA extraction kit (Invitrogen™). The quality of total RNA was assessed on a BioAnalyzer 2100 (Agilent Technologies, Santa Clara, CA, USA) to ensure a RIN > 9 for all samples. Mature mRNA was extracted from 1ug of total RNA and cDNA libraries were constructed using a TruSeq kit (Illumina).

Quality control of libraries was assessed on a BioAnalyzer and quantification measured using qPCR. *NEBNext* Multiplex adaptors were added via ligation, and the cDNA libraries were sequenced at Genome Quebec, McGill University. Three libraries were multiplex sequenced on one lane of Illumina at a concentration of 8 pM per cDNA library using HiSeq2500 transcriptome sequencing to generate 125bp paired-end reads. This resulted in approximately 80 million reads for each transcriptome.

*Tetrapygus niger*. Adult specimens were collected and transported alive to the University of Concepcion. Tissue samples (female gonad and tube feet) from one individual were finely chopped with a scalpel, and total RNA was extracted immediately using TriReagent (Sigma), following the manufacturer’s instructions. Total RNA concentration was estimated using QuantiFluor® RNA System (111.84-339.04 ng/µL) and quality was assessed using Agilent RNA 6000 Nano kit with an Agilent 2100 TapeStation (RIN: 5.4-8.4). cDNA libraries were prepared at Genoma Mayor SpA (Chile) using TruSeq Stranded mRNA Kit (Illumina) and sequenced on a multiplexed run using Illumina HiSeq 4000 platform with 150 paired-end reads, resulting in 54.1 M reads for gonad female and 52.7 M reads for tube feet, respectively. These two transcriptomic datasets were combined before performing all steps of bioinformatic processing.

*Eucidaris metularia*. A single specimen was preserved in RNA*later* after being starved overnight and kept for 1 day at 4°C before long-term storage at -80°C. Total RNA was extracted using Direct-zol RNA Miniprep Kit (with in-column DNase treatment; Zymo Research) from Trizol. mRNA was isolated with Dynabeads mRNA Direct Micro Kit (Invitrogen). RNA concentration was estimated using Qubit RNA broad range assay kit, and quality was assessed using RNA ScreenTape with an Agilent 4200 TapeStation. Values were used to customize downstream protocols following manufacturers’ instructions. Library preparation was performed with KAPA-Stranded RNA-Seq kits, targeting an insert size in the range of 200–300 base pairs. The quality and concentration of libraries were assessed using DNA ScreenTape.

The library was sequenced in a multiplexed pair-end runs using Illumina HiSeq 4000 with 7 other libraries in the same lane (all previously published in [4]). In order to minimize read crossover, 10 bp sequence tags designed to be robust to indel and substitution errors were employed [113].

The assembled transcriptome was filtered from cross-contaminant reads product of multiplexed sequencing using CroCo v. 1.1 [91]. Seven other transcriptomes sequenced together were also employed, and results of this step are reported in Fig. S1 of Mongiardino Koch & Thompson [3].

### Fossil constraints

All clade names used below refer to the corresponding crown groups.

<u>Ambulacraria (Echinodermata-Hemichordata divergence)</u> – **Younger bound**: 518 Ma, Middle Atdabanian (age 3, Series 3 of Cambrian). **Older bound**: 636.1 Ma, Lantian Biota, Ediacaran. **Reference:** [118]. **Notes:** The root of our tree is constrained with a younger bound concurrent with the earliest occurrence of stereom in the fossil record (see below). The older bound is based on the maximum age of the Lantian Biota, a Lagerstätte lacking any trace of eumetazoan fossils.

<u>Echinodermata (Pelmatozoan-Eleutherozoan divergence)</u> – **Younger bound**: Unconstrained. **Older bound:** 515.5 Ma, Middle Atdabanian (age 3, Series 3 of Cambrian). **Reference**: [118]. **Notes:** This divergence (which represents the divergence of crown group echinoderms), has an older bound based upon the earliest occurrence of stereom in the fossil record. Stereom constitutes an echinoderm synapomorphy readily recognizable in the fossil record due to its unique mesh-like structure [43]. The first records of stereom point to a sudden and global appearance within the middle Atdabanian [42]. Note that this is also used as younger bound for the divergence between echinoderms and hemichordates.

<u>Echinozoa (Echinoidea-Holothuroidea divergence)</u> – **Younger bound:** 469.96 Ma, Top of

*Pseudoclamacograptus decorates* graptolite zone. **Older bound:** 461.95 Ma, Top Floian. **Reference:** [119, 120]. **Notes:** The oldest crown eleutherozoan fossils are disarticulated holothurian calcareous ring elements which are known from the Red *Orthoceras* limestone of Sweden. This falls within the *P. decorates* graptolite zone.

<u>Asterozoa (Ophiuroidea-Asteroidea divergence)</u> – **Younger bound**: 521 Ma, Top of *Psigraptus jacksoni*one. **Older bound**: 480.55 Ma, Base of Series 2 Cambrian Period. **Reference**: [119, 120]. **Taxon**: *Maydenia roadsidensis.* **Notes**: The divergence of asterozoan classes is calibrated based upon the stratigraphic occurrence of *Maydenia roadsidensis*, the oldest asterozoan, which is a stem group ophiuroid [121] and occurs in the *Psigraptus jacksoni* Zone of the Floian.

<u>Asteroidea</u> – **Younger bound:** 239.10 Ma, Top Fassanian, Ladinian, Mo3, *nodosus* biozone, Triassic.

**Older bound**: 252.16 Ma, P-T Boundary. **Reference**: [122]. **Taxon**: *Trichasteropsis weissmanni*. **Notes**: Fossil evidence suggests the divergence of the asterozoan crown group took place sometime in the Early or Middle Triassic [122]. Based upon its phylogenetic position and stratigraphic occurrence as the oldest crown group asteroid, we use the forcipulatacean *T. weissmanni* from the Middle Triassic Muschelkalk as the soft bound on this divergence.

<u>Crinoidea</u> – **Younger bound**: 247.06 Ma, Top Spathian (Paris Biota, Lower Shale unit, Thaynes Group, Spathian, Triassic). **Older bound**: Unconstrained. **Reference**: [123]. **Taxon**: *Holocrinus*. **Notes**: Crown group Crinoidea is difficult to define, though a rapid post-Palaeozoic morphological diversification is supported by fossil evidence. We use *Holocrinus* from the Early Triassic of the Thaynes group to calibrate the younger bound on the divergence of the crown group (i.e., the split between Isocrinida and all other extant crinoids).

<u>Ophiuroidea (Euryophiurida-Ophintegrida divergence)</u> – **Younger bound**: 247.06 Ma, Top Spathian (Paris Biota, Lower Shale unit, Thaynes Group, Spathian, Triassic). **Older bound**: Unconstrained. **Reference**: [124]. **Taxon**: *Shoshonura brayardi*. **Notes**: *S. brayardi* from the Spathian Thaynes Group of the Western USA is a member of the crown group ophiuroid clade Ophiodermatina. It is thus the currently oldest described member of the ophiuroid crown group, setting its minimum age of divergence.

<u>Pneumonophora (Holothuriida-Neoholothuriida divergence)</u> – **Younger bound**: 259.8 Ma, S*pinosus* zone of Early Ladinian (used base Ladinian). **Older bound**: 241.5 Ma, Base Wuchiapingian. **Reference**: [46, 119]. **Notes**: The oldest (undescribed) holothuriid calcareous ring ossicles are from the *Spinosus* zone of the Ladinian of Germany and calibrate its divergence from neoholothuriids.

<u>Echinoidea (Cidaroidea-Euechinoidea divergence)</u> – **Younger bound**: 298.9 Ma, Base Carnian. **Older bound**: 237 Ma, Base Permian. **Reference**: [125]. **Taxon**: *Triassicidaris? ampezzana*. **Notes**: The recent phylogenetic analysis of Mongiardino Koch and Thompson [3] found many traditional early crown group echinoids as members of the stem group. Given this result, *Triassicidaris? ampezzana*, a cidaroid which is known from the St. Cassian beds of Italy, becomes the current oldest crown group echinoid.

<u>Pedinoida-Aspidodiadematoida</u> – **Younger bound**: 209.46 Ma, Top of Norian. **Older bound**: 252.16 Ma, P-T boundary. **Reference**: See discussion in supplement of Thompson et al. [13]. **Taxon**: *Diademopsis* ex. gr. *heberti*. **Notes**: The oldest euechinoid, *Diademopsis* ex. gr. *heberti*, which is a pedinoid, is known from the Norian of Peru. This fossil calibrates the younger bound on the pedinoid-aspidodiadematoid divergence.

<u>Carinacea (Echinacea-Irregularia divergence)</u> – **Younger bound**: 234.5 Ma, Top Sinemurian. **Older bound**: 190.8 Ma, Top Julian 1 ammonoid zone of Norian. **Reference**: Li et al. [126]. **Taxon**: *Jesionekechinus*. **Notes**: The divergence of Carinacea is calibrated based upon the oldest irregular echinoid, *Jesionekechinus*, which occurs above the Sinemurian of New York Canyon, Nevada. For a more detailed discussion of this divergence see the supplement of Thompson et al. [13].

<u>Salenioida-(Camarodonta+Stomopneustoida)</u> divergence– **Younger bound**: 228.35 Ma, Base Hettangian.

**Older bound**: 201.3 Ma, Top Carnian. **Reference**: [14]. **Taxon**: *Acrosalenia chartroni*. **Notes**: The stem group salenioid *Acrosalenia chartroni* from the Hettangian of France is the earliest representative of Echinacea. Given the topology recovered by our analyses, this fossil was used to calibrate the divergence between salenioids and the clade composed of stomopneustoids and camarodonts.

<u>Stomopneustoida-Camarodonta divergence</u> – **Younger bound**: 201.3 Ma, Top Pliensbachian. **Older bound**: 182.7 Ma, Base Jurassic. **Reference**: See discussion in supplement of Thompson et al. [13]. **Taxon**: Stomechinids from Morocco like *Diplechinus hebbriensis* [127]. **Notes**: The oldest stomechinids, which are stem group stomopneustoids, are from the Early Jurassic of Morocco.

<u>Neognathostomata-Atelostomata divergence</u> – **Younger bound**: 234.5 Ma, Base of Toarcian. **Older bound**: 182.7 Ma, St. Cassian Beds, bottom of Julian 2 ammonoid Zone. **Reference**: [14, 126]. **Taxon**: Younger bound is set by *Galeropygus sublaevis* (older bound is relatively uninformative). **Notes**: The base of the Toarcian (which contains the oldest neognathostomate *G. sublaevis*) is used as the lower bound on the divergence between the neognathostomata and all other crown irregular echinoids. Note that depending on the position of the unsampled echinoneoids, this split might also correspond to the origin of crown group Irregularia (see recent topologies recovered by Lin et al. [36] and Mongiardino Koch & Thompson [3]).

<u>Atelostomata (Holasteroida-Spatangoida divergence)</u> – **Younger bound**: 137.68 Ma, Bottom of *Campylotoxus* zone in Valanginian. **Older bound**: 201.3 Ma, Base of Jurassic. **Reference**: [128]. **Taxon**: Younger bound is based on *Toxaster* (older bound is loosely uninformative). **Notes**: The toxasterids are stem group spatangoids that calibrate the divergence between spatangoids and holasteroids. The oldest *Toxaster* are from the *Campylotoxus* zone in the Valanginian, and set the younger bound on the divergence.

<u>Echinolampadacea (Cassiduloida-Scutelloida divergence)</u> – **Younger bound**: 113 Ma, Base Albian. **Older bound**: 145 Ma, Base Cretaceous. **Reference**: [129]. **Taxon**: Younger bound is based on *Eurypetalum rancheriana* (older bound is loosely uninformative). **Notes**: The oldest member of Echinolampadacea is the faujasiid *Eurypetalum rancheriana* (originally placed in the genus *Faujasia* and later transferred, see [28, 29]) from the Late Albian of Colombia.

<u>Scutelloida (Scutelliformes-Laganiformes divergence)</u> – **Younger bound**: 56.0 Ma, Bottom of Eocene.

**Older bound**: Unconstrained. **Reference**: [14, 130]. **Taxon**: Younger bound is based on *Eoscutum doncieuxi*. **Notes**: We use the base of the Eocene as the younger bound on the divergence between scutelliforms and laganiforms. This is based upon the occurrence of *E. doncieuxi*, the oldest scutelliform, which is known from the Lower Eocene. The older bound is left unconstrained in order to account for the current uncertainty in the origin of the clade (i.e., allow for Mesozoic ages).

*<u>Leodia</u>*<u>-(*Encope*+*Mellita*) divergence</u> – **Younger bound**: 23.03 Ma, Base Miocene. **Older bound**: 33.9 Ma, Base of Oligocene. **Reference**: [131, 132]. **Taxon**: Younger bound is based on *Encope michoacanensis* (older bound is loosely uninformative). **Notes**: This node is constrained using the oldest representative of these three genera of mellitids.

<u>Clypeasteroida</u> – **Younger bound**: 47.8 Ma, Base Lutetian. **Older bound**: Unconstrained. **Reference**: [133, 134]. **Taxon**: Younger bound is based on *Clypeaster calzadai* and *Clypeaster moianensis*. **Notes**: The oldest known fossil species of Clypeasteroida (the clypeasterids *C. calzadai* and *C. moianensis*) are from the middle Eocene, upper Lutetian (Biarritzian) of Cataluña, Spain. The older bound is left unconstrained in order to account for the current uncertainty in the origin of the clade (i.e., allow for Mesozoic ages).

<u>Strongylocentrotidae-Toxopneustidae divergence</u> – **Younger bound**: 44.4 Ma, Eocene Planktonic Zones E9-E12. **Older bound**: 56.0 Ma, Base Eocene. **Reference**: [135, 136]. **Taxon**: Younger bound is based on *Lytechinus axiologus* (older bound is loosely uninformative). **Notes**: The oldest member of the clade comprising toxopneustids and strongylocentrotids is the toxopneustid *Lytechinus axiologus.* This taxon is known from the Eocene Yellow Limestone Group of Jamaica. The Yellow Limestone group represents the Eocene Planktonic Zones E9-E12.

## Supplementary Figures and Tables

**Figure S1:**
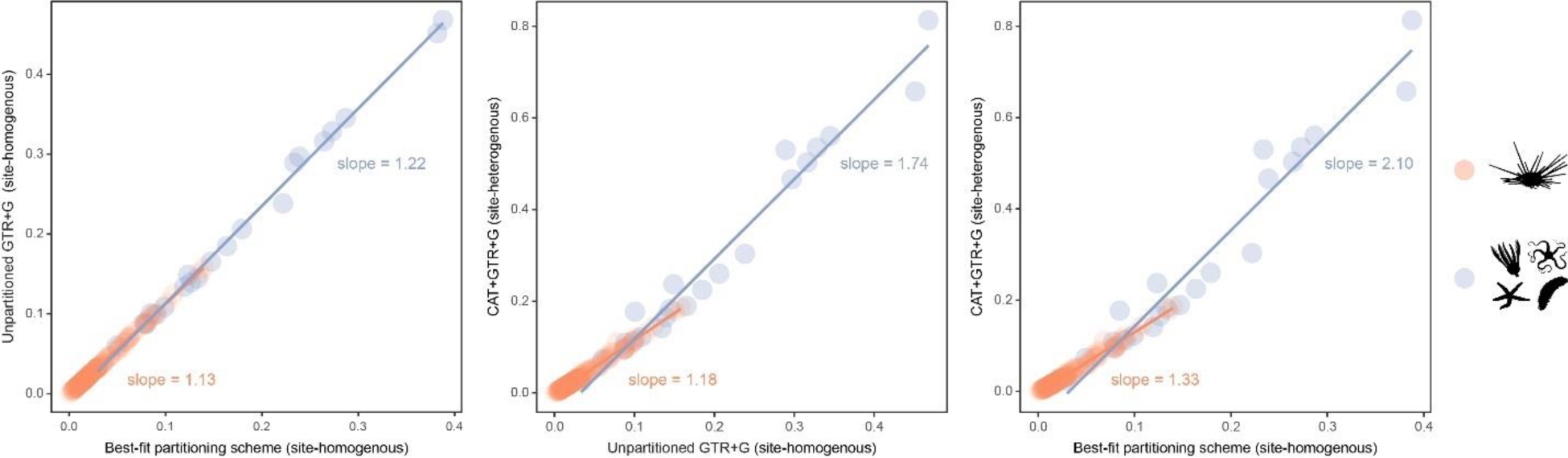
Estimated branch lengths across different models of molecular evolution. Different site-homogeneous models (left) infer similar levels of divergence, and the choice between them induces little distortion in the general tree structure. Site-heterogeneous models on the other hand, not only infer a larger degree of divergence between terminals relative to site-homogeneous ones (center and left), but they also distort the tree (i.e., impose a non-isometric stretching), with branch lengths connecting outgroup taxa expanding much more than those within the ingroup clade.

**Figure S2:**
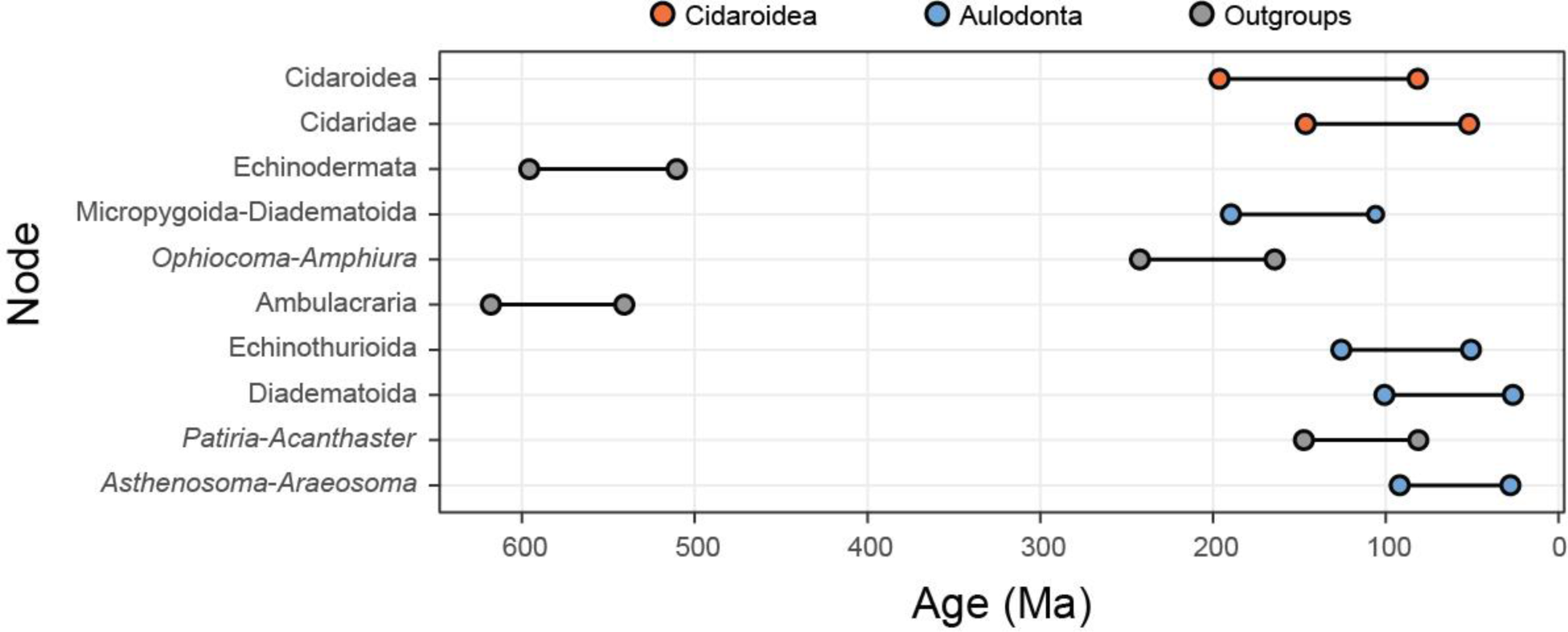
The ten most sensitive node dates are found within Cidaroidea, Aulodonta, and among outgroup nodes. For each, the range shown spans the interval between the minimum and maximum ages found among the consensus topologies of all time-calibrated runs.

**Figure S3:**
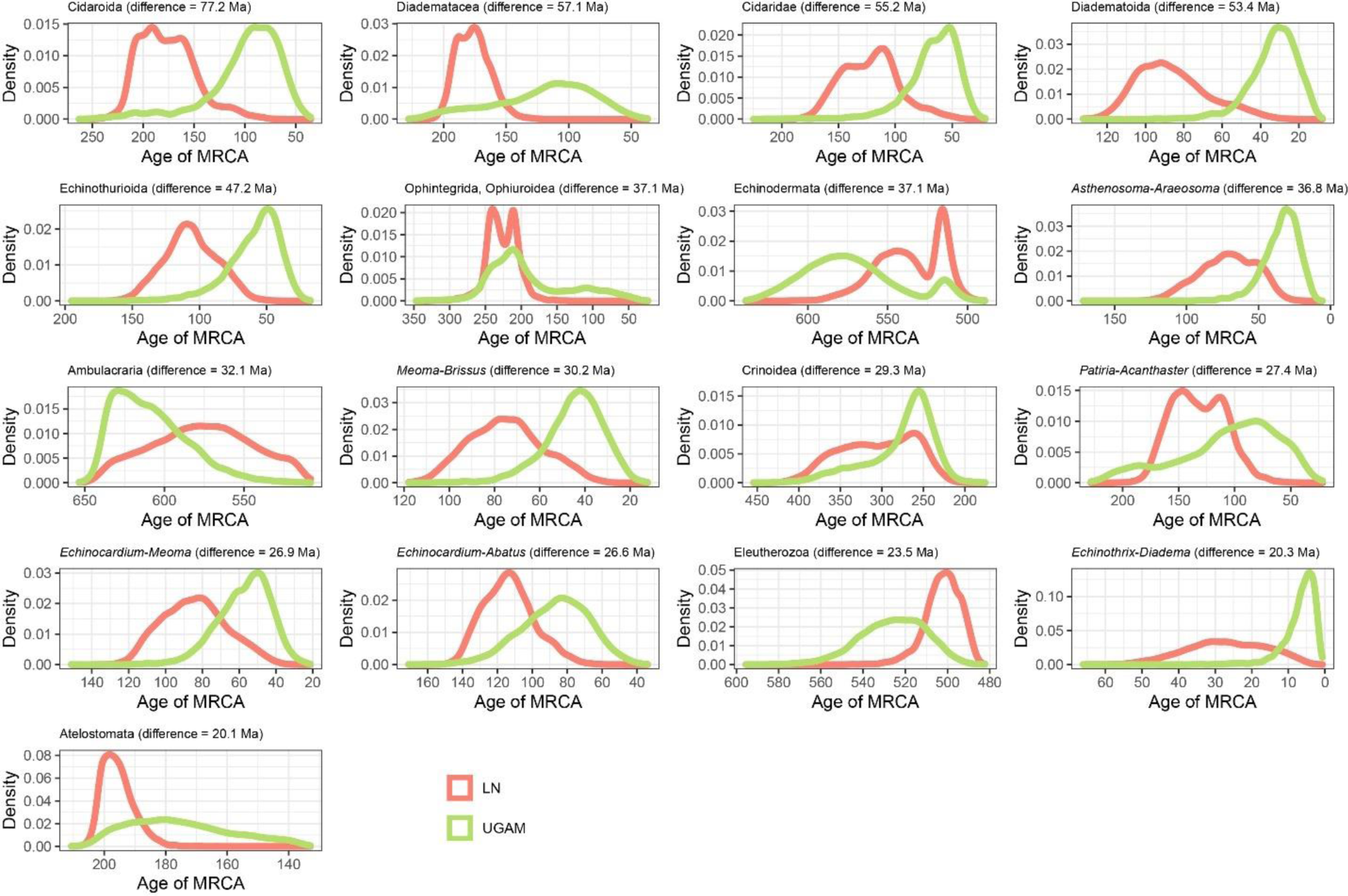
Distribution of posterior probabilities for node ages that show an average difference larger than 20 Ma depending on the choice of clock prior.

**Figure S4:**
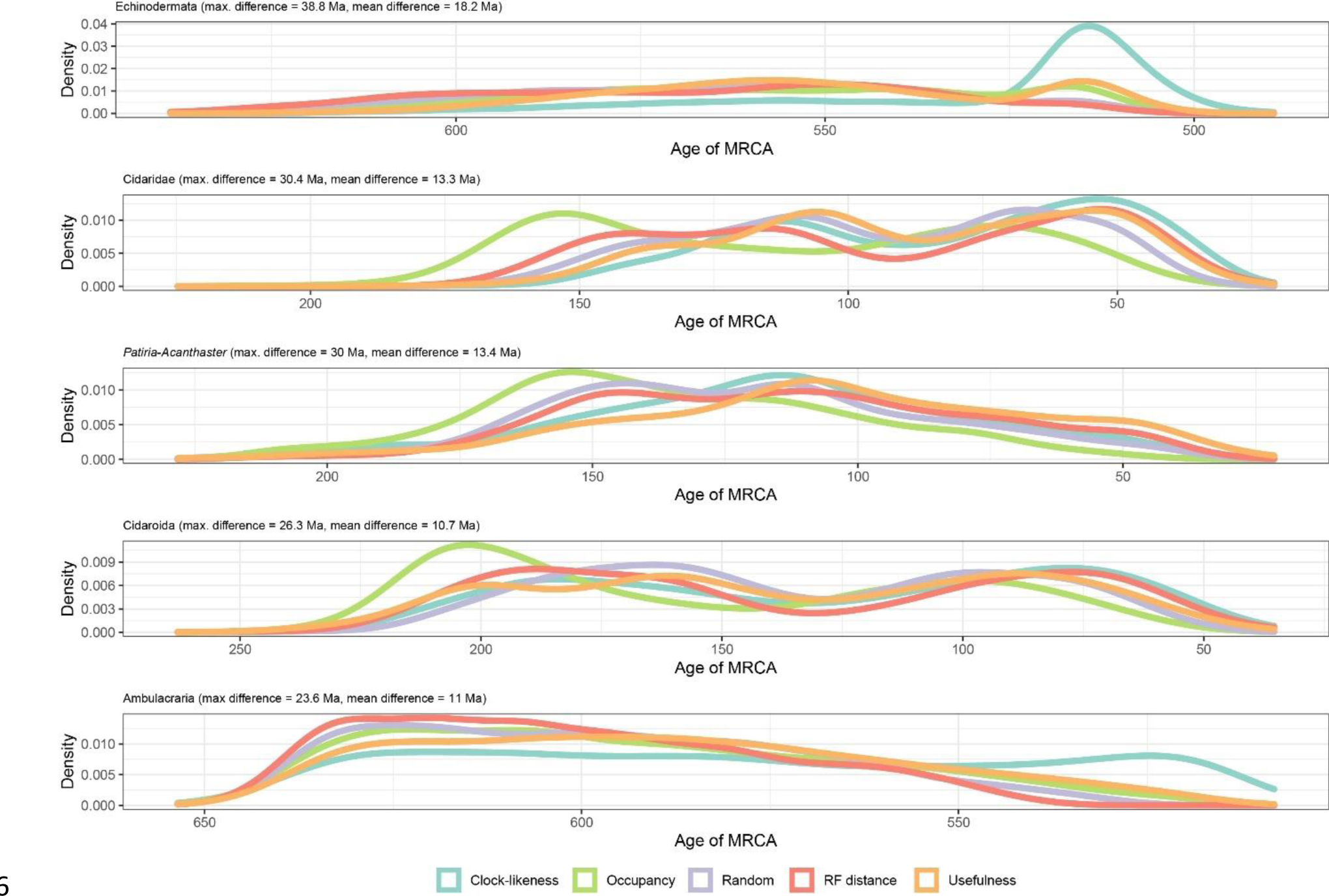
Distribution of posterior probabilities for node ages that show a maximum difference larger than 20 Ma depending on the choice of clock prior. The largest differences can be seen in the relatively younger ages of Ambulacraria and Echinodermata when using clock-like genes, and in the relatively older ages for some nodes within cidaroids and starfish when using loci with high occupancy. Other sampling criteria largely agree on inferred node ages, as can also be seen in Fig. 3C as short distances between their centroids in chronospace.

**Figure S5:**
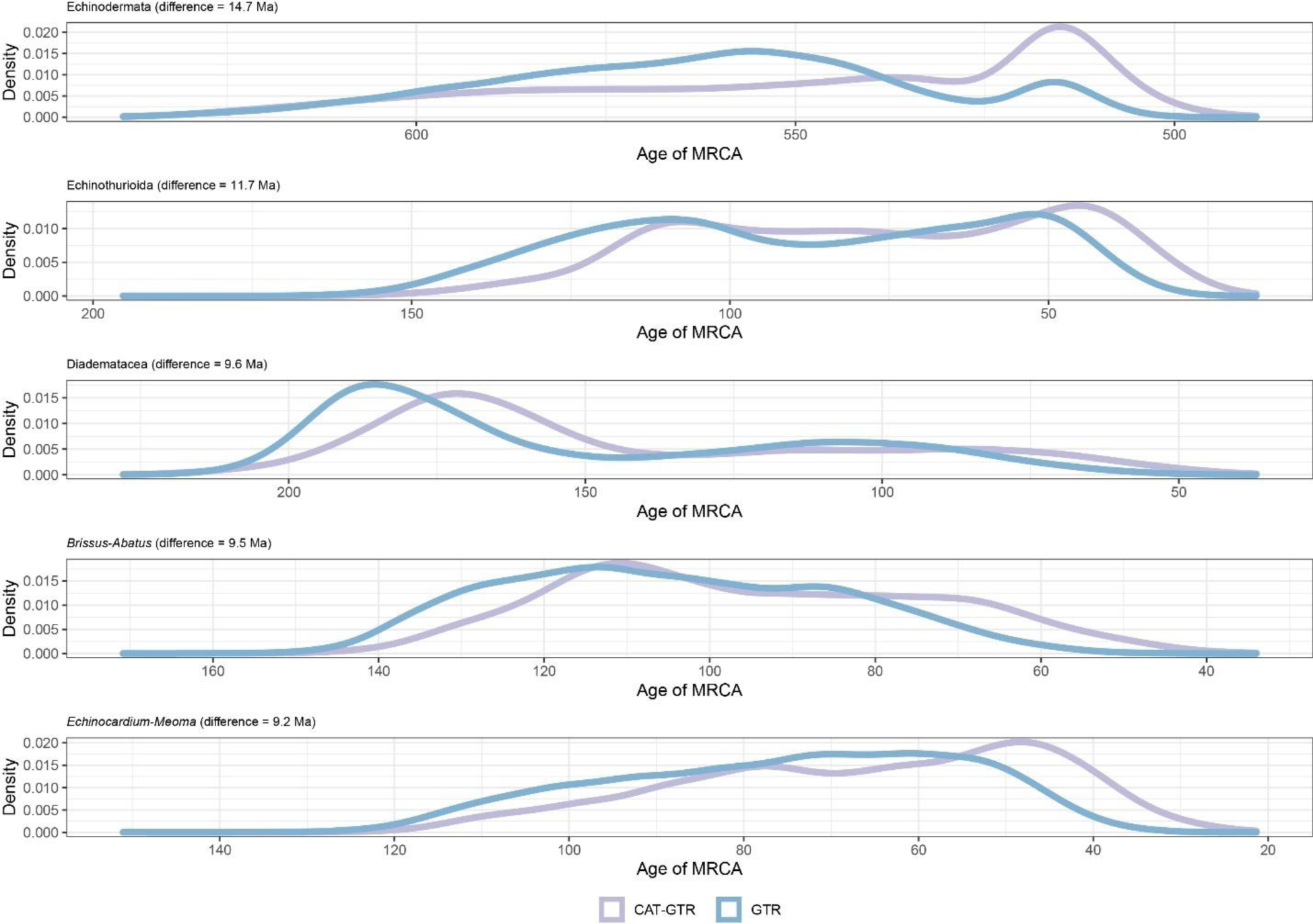
Distribution of posterior probabilities for node ages that are the most affected by the choice of model of molecular evolution. No node showed average differences larger than 20 Ma, so those with the largest effect are shown.

**Figure S6:**
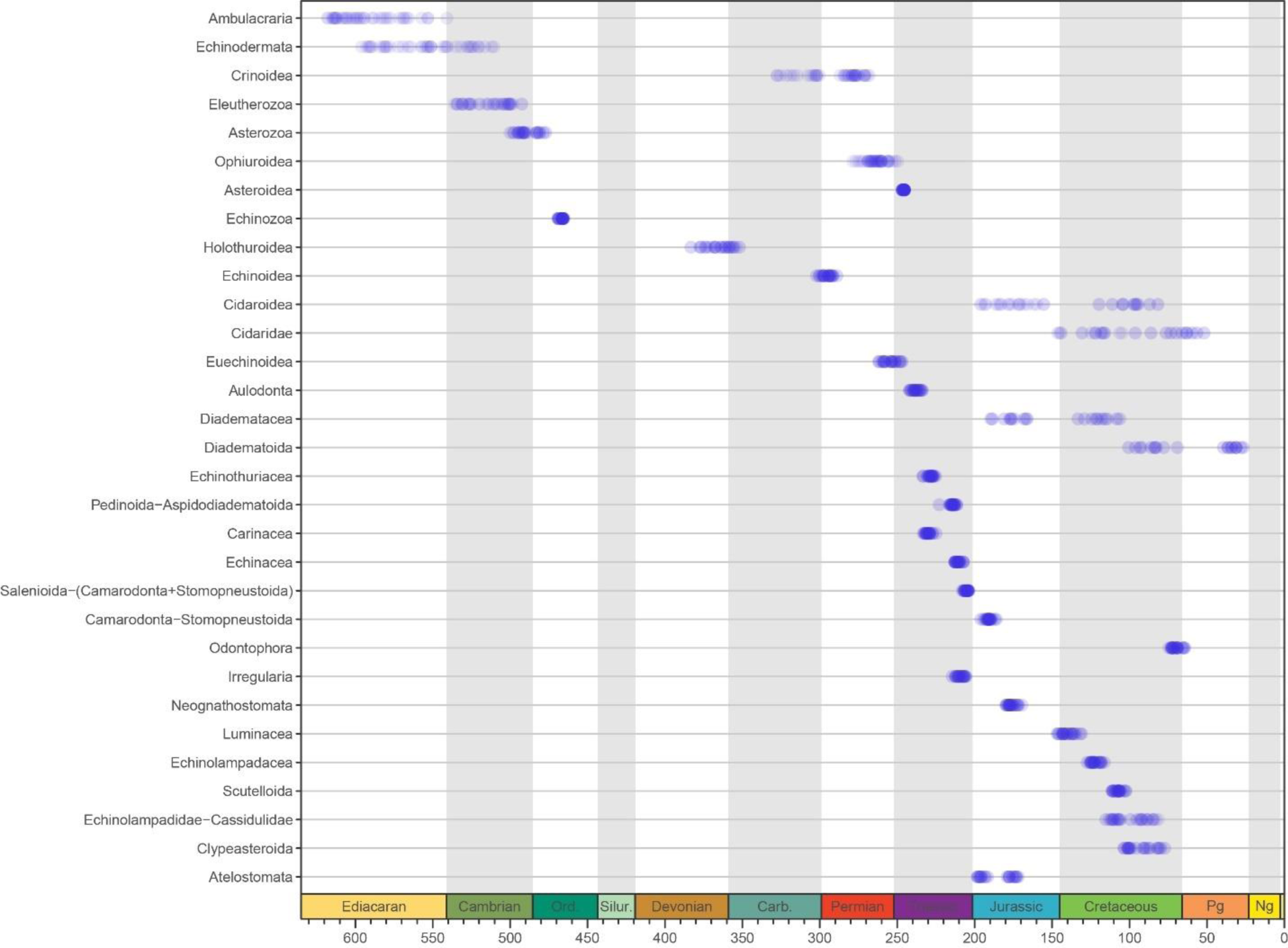
Median ages for selected clades across the consensus trees of the 40 time-calibrated experiments performed (20 analyses x 2 runs each).

**Figure S7:**
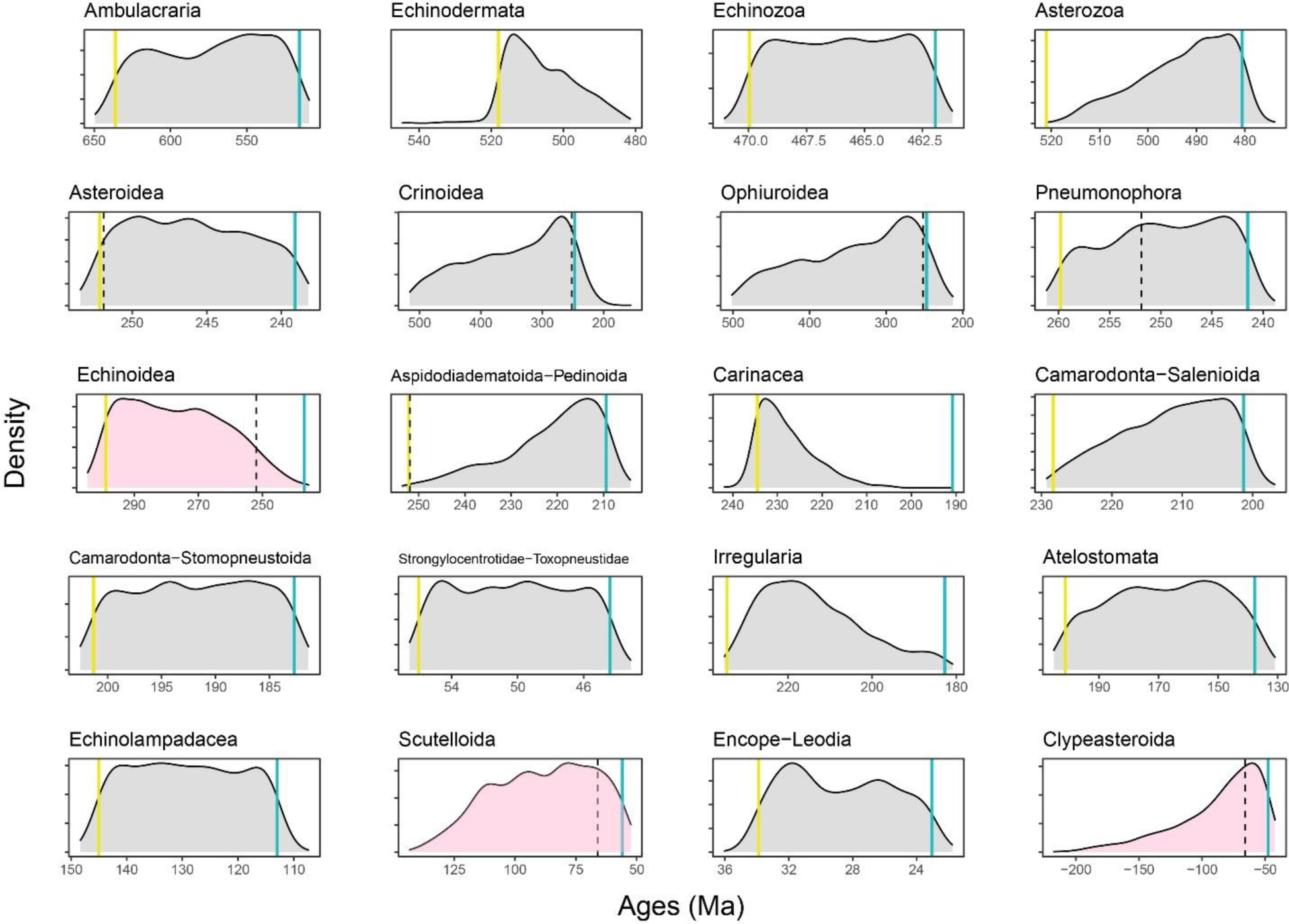
Prior distributions of all constrained nodes. Five hundred replicates were sampled from the joint prior, showing appropriately broad distributions of node ages. Blue lines show minima and yellow ones maxima (when enforced); dotted lines show the age of the Permian-Triassic (251.9 Ma) and Cretaceous-Paleogene (66 Ma) mass extinction events. Nodes whose ages are of special interest (Echinoidea, Scutelloida, and Clypeasteroida) are shown in pink, revealing large prior probabilities of the divergence occurring at either side of mass extinction events.

**Figure S8:**
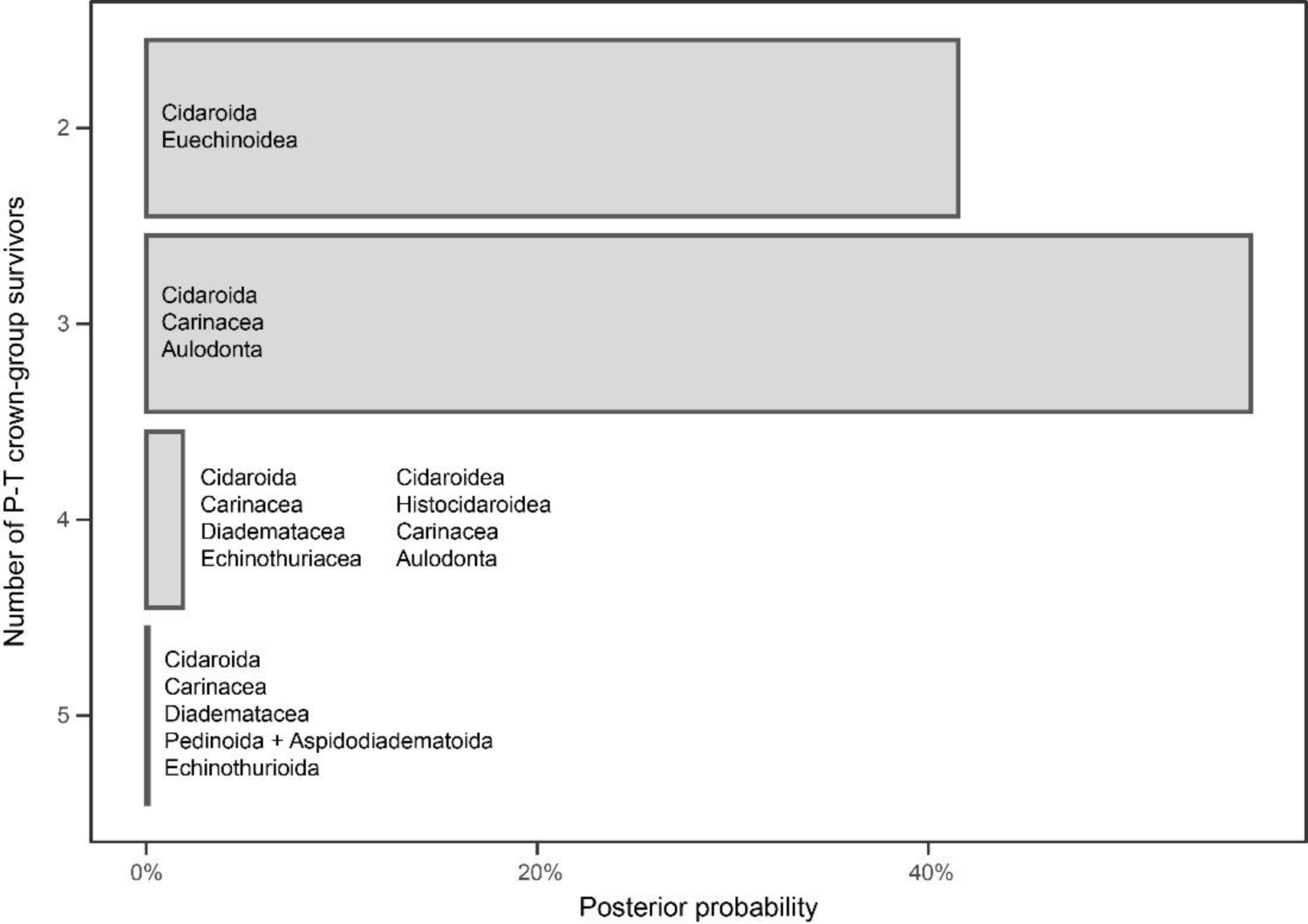
Number of lineages inferred to have crossed the Permian-Triassic (P-T) boundary. The probabilities of each scenario are estimated from the inferred divergence times of the 10,000 chronograms sampled across all of the analyses performed (250 for the two runs of each combination of sampled loci, model of molecular evolution, and clock prior). The probability of three or more crown group lineages surviving the Permian-Triassic extinction is 58.47%.

**Figure S9:**
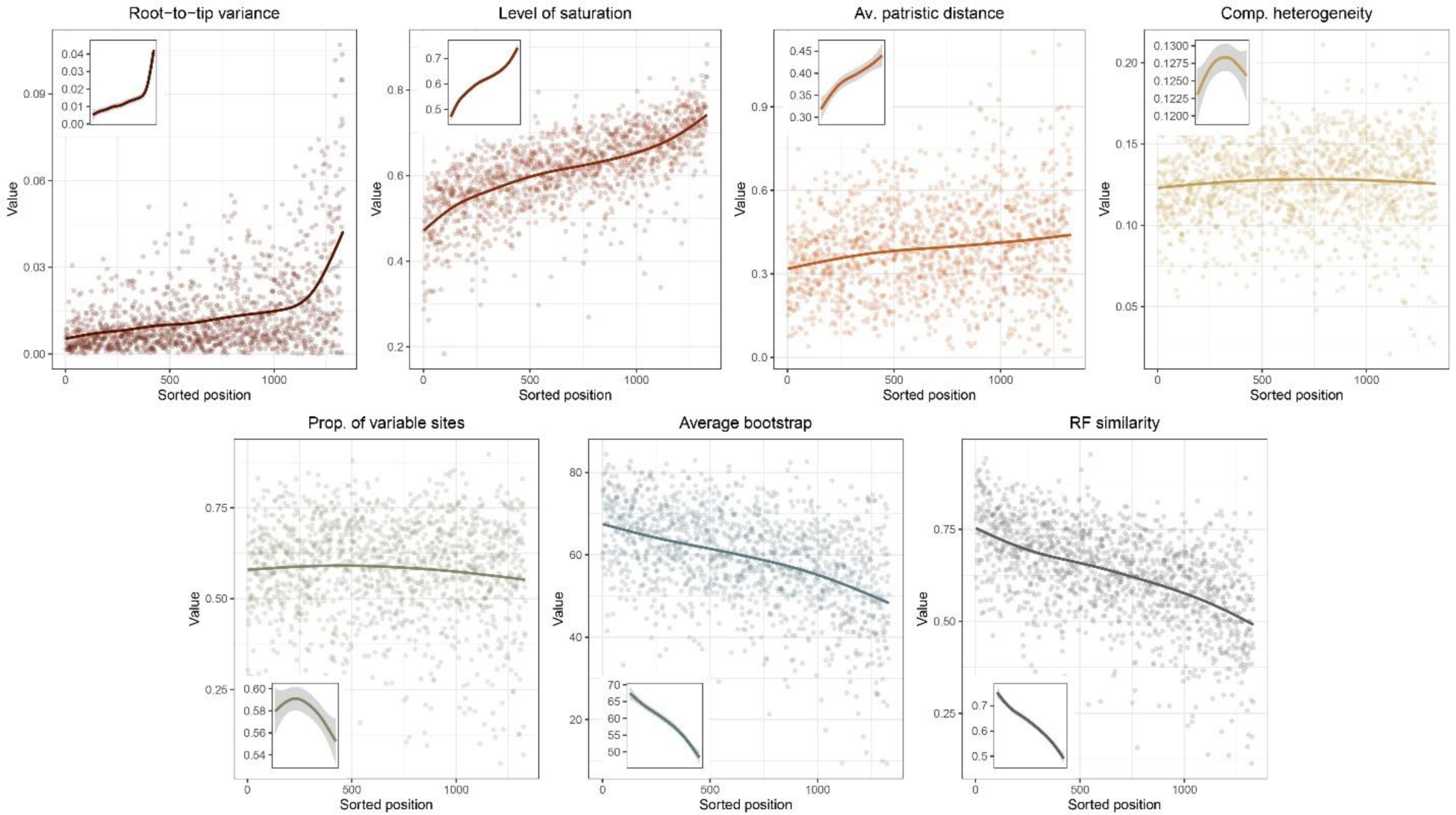
Ordering of loci enforced using *genesortR* [37] and its relationship to the seven gene properties employed. High ranking loci (i.e., the most phylogenetically useful) show low root-to-tip variances (or high clock-likeness), low saturation and compositional heterogeneity, as well as high average bootstrap and Robinson-Foulds similarity to a target topology (in this case, with the contentious relationship among major lineages of Echinacea collapsed).

**Figure S10:**
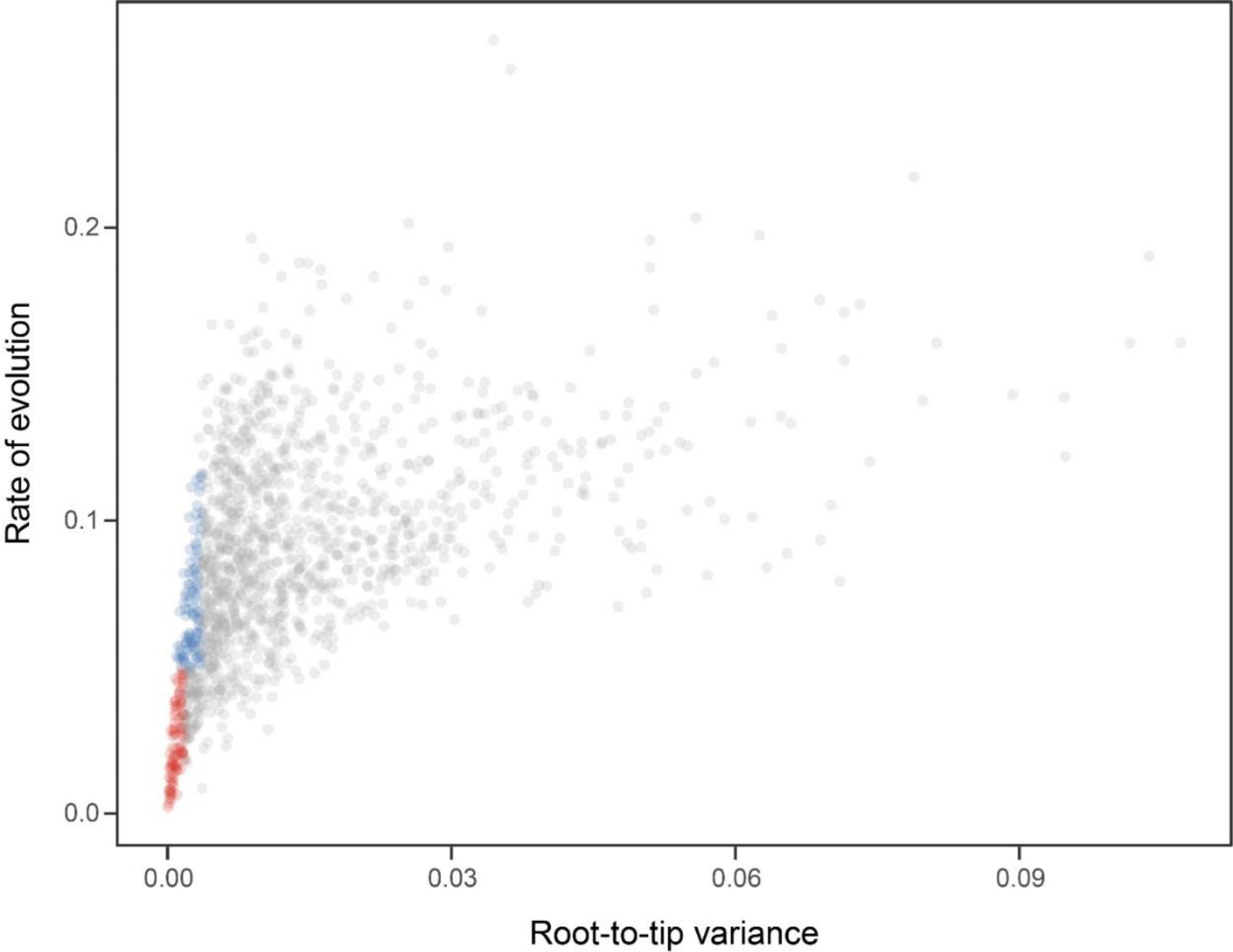
Relationship between the root-to-tip variance (a proxy for the clock-likeness of loci) and the rate of evolution (estimated as the total tree length divided by the number of terminals). The most clock-like loci (shown in red), which are often favored for the inference of divergence times (e.g., [62, 72]), are among the most highly conserved and can provide little information for constraining node ages [37]. Highly clock-like genes with a higher information content were used instead by choosing the loci with the lowest root-to-tip variance from among those that were within 1 standard deviation from the mean evolutionary rate (shown in blue).

**Figure S11:**
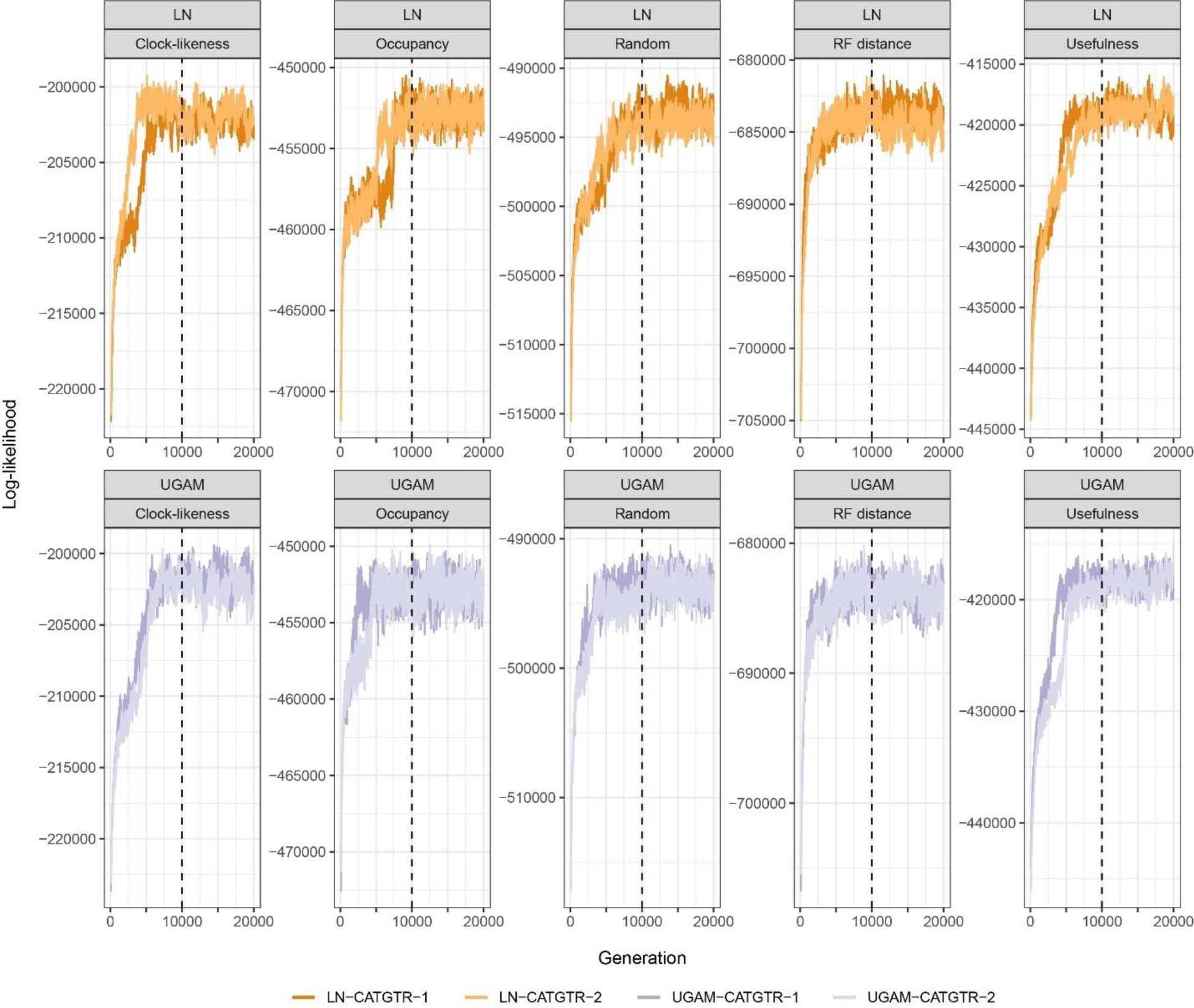
Trace plots of the log-likelihood of different time-calibration runs. All runs show evidence of reaching convergence and stationarity before our imposed burn-in fraction of 10,000 generations. For simplicity, only runs under the CAT+GTR+G model are plotted. Those run under GTR+G converged much faster.

**Table S1:**
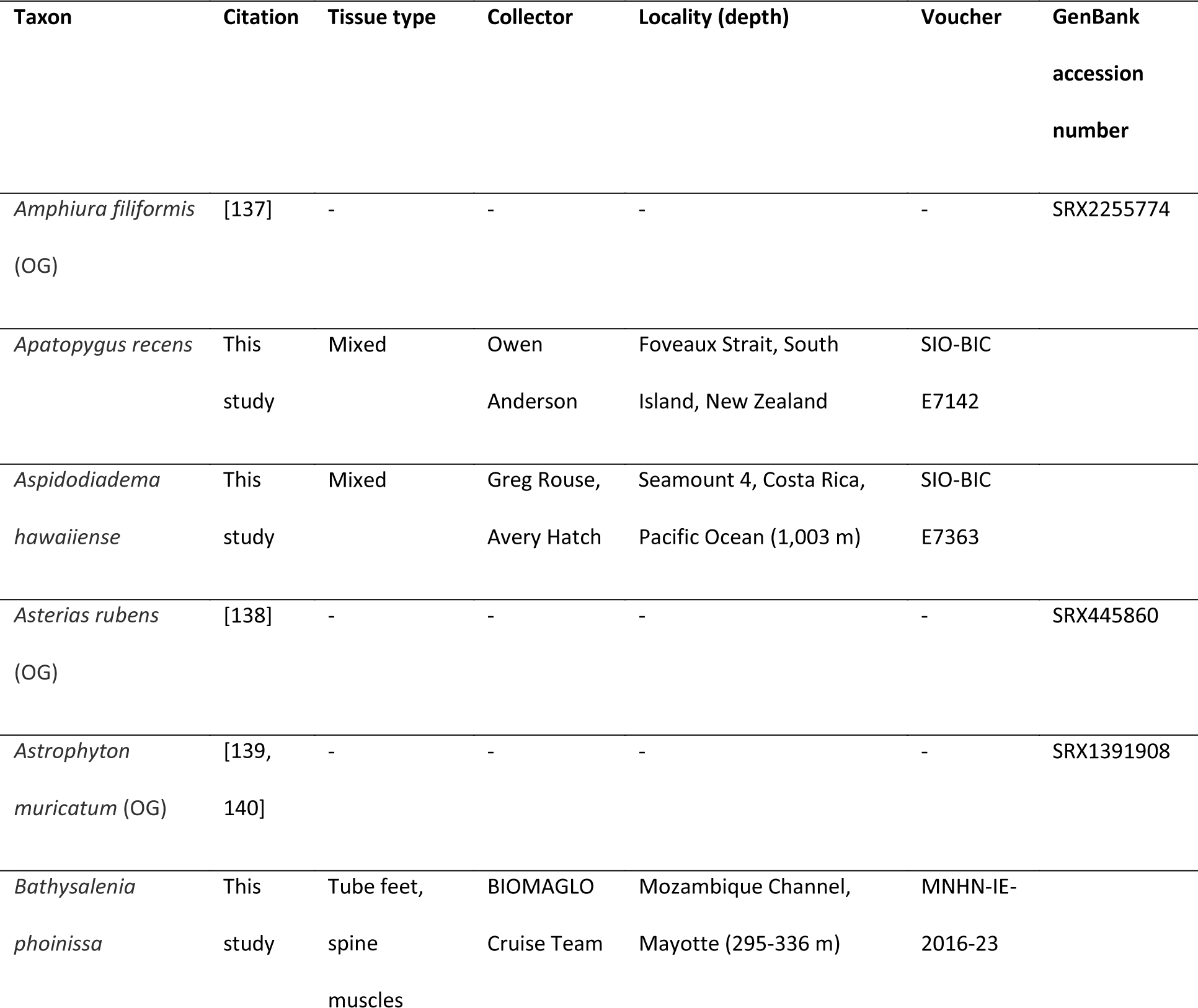
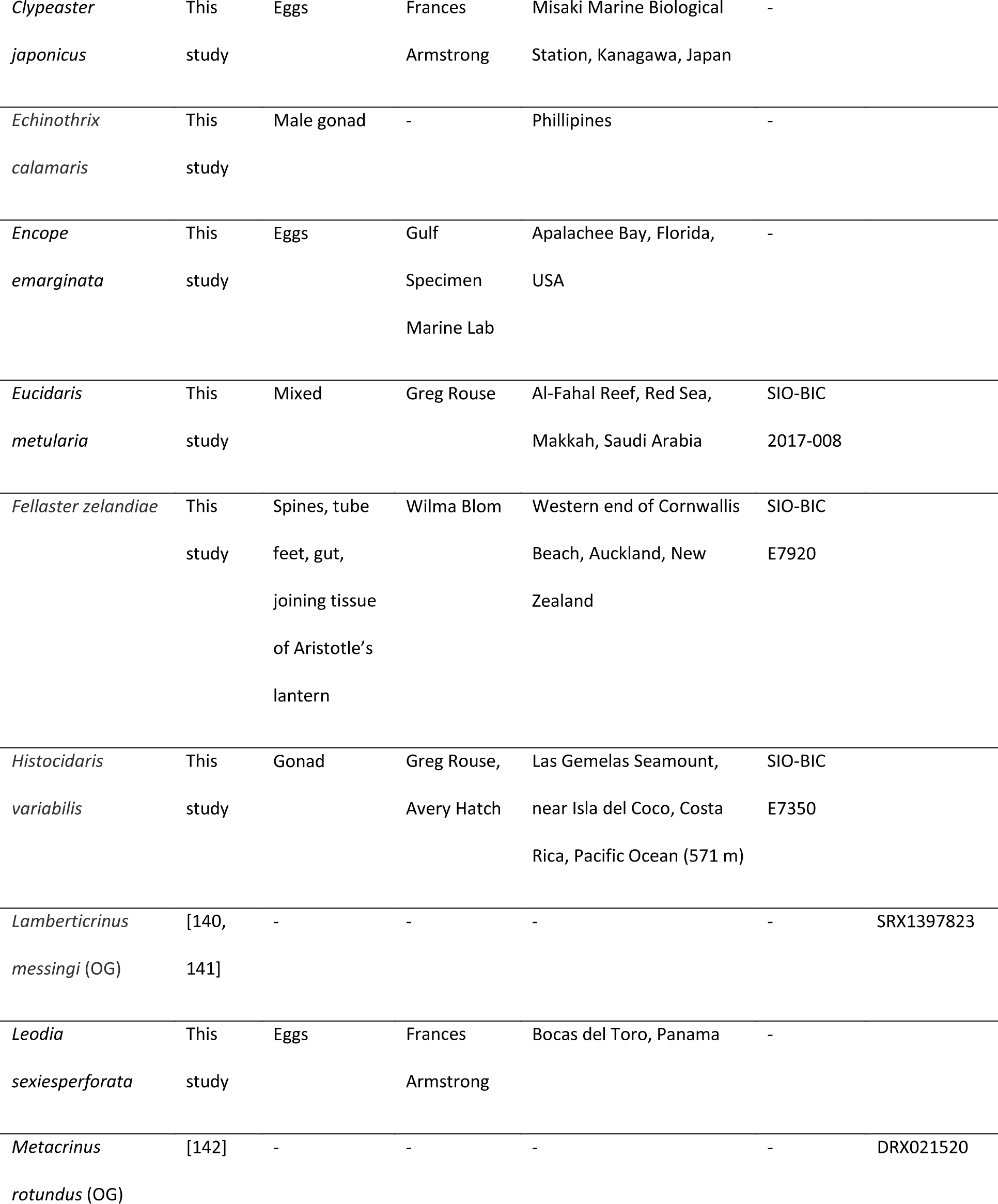
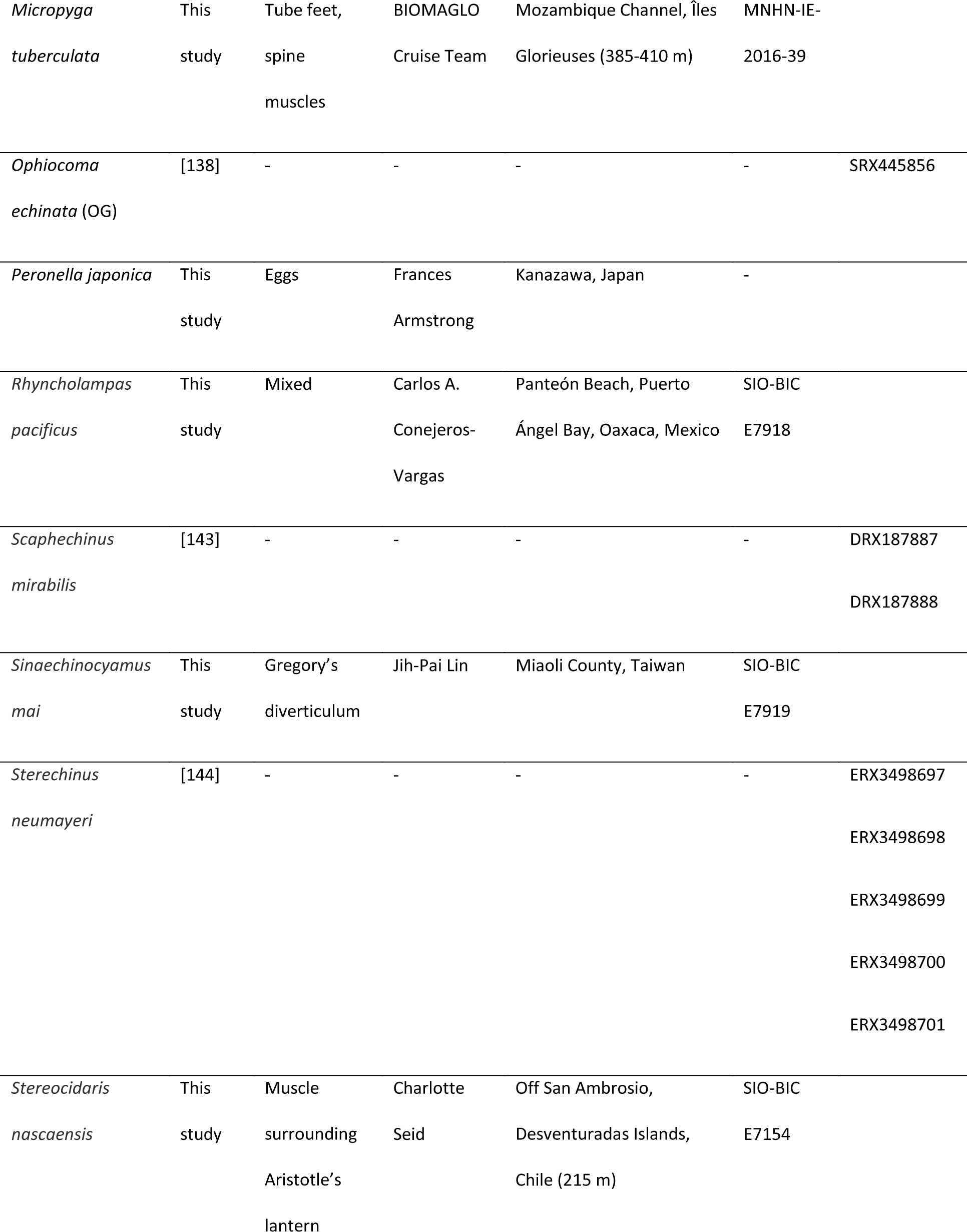
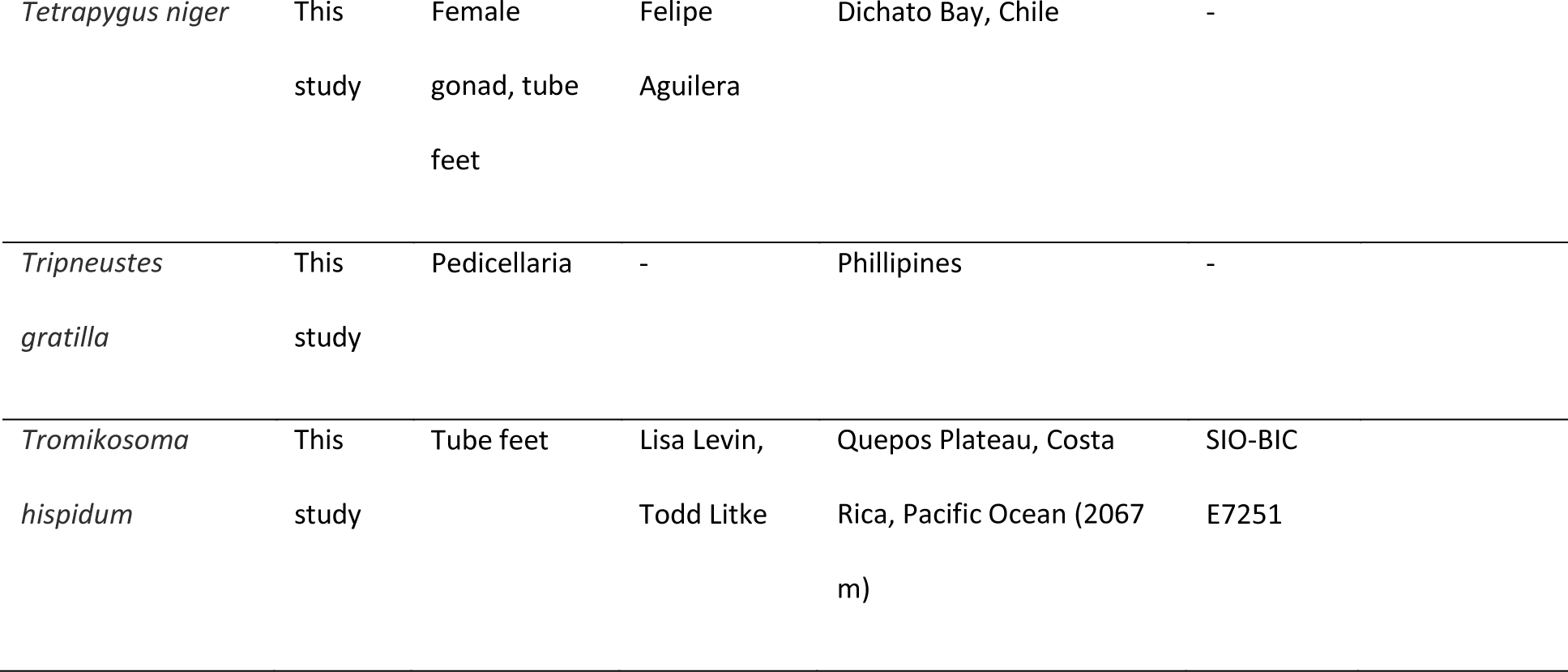
Transcriptomic/genomic datasets added in this study relative to the taxon sampling of Mongiardino Koch et al. [4] and Mongiardino Koch & Thompson [3]. Taxa with citations were taken from the literature, and details can be found in the corresponding studies and associated NCBI BioProjects. Taxa are shown in alphabetical order; those identified with “OG” are outgroup taxa (i.e., non-echinoids). Voucher specimens are deposited at the Benthic Invertebrate Collection, Scripps Institution of Oceanography (SIO-BIC), and the Echinoderm Collection, Muséum National d’Histoire Naturelle (MNHN- IE). If multiple accession numbers are shown for a given taxon, these datasets were merged before assembly. Similar details for all other specimens can be found in [3, 4].

**Table S2:** Details of molecular datasets and supermatrix. Statistics for raw reads and assemblies are shown for datasets incorporated in this study relative to the sampling of [3, 4] (where similar statistics can be found for the other datasets). Taxa are shown in alphabetical order; those identified with “OG” are outgroup taxa (i.e., non-echinoids). Novel datasets correspond to Illumina pair-end transcriptomes, except for two draft genomes (*Bathysalenia phoinissa* and *Micropyga tuberculata*). Mean insert size is expressed in number of base pairs. For transcriptomes, read pairs (initial) shows numbers input into Agalma [84], (i.e., after processing with Trimmomatic [83] or UrQt [86]), while read pairs (retained) shows those that passed the Agalma curation checks (including ribosomal, adapter, quality, and compositional filters), and represent the final number of read pairs used for assembly. For genomes, see information in the bioinformatic details above. Assemblies were further sanitized with either alien_index [92] alone or in combination with CroCo [91] (see bioinformatic details), and the number of assembled transcripts/contigs shows the size of datasets after these curation steps. The number of loci shows the occupancy of terminals in the supermatrix output by Agalma (1,346 loci at 70% occupancy), after which loci were further removed by TreeShrink [97], resulting in the final occupancy scores.

| Species | Mean<br>insert size | Read pairs<br>(initial) | Read pairs<br>(retained) | Assembled<br>transcripts/contigs | Number<br>of loci | Removed by<br>TreeShrink | Final<br>occupancy |
| --- | --- | --- | --- | --- | --- | --- | --- |
| *Abatus agassizii* | - | - | - | - | 522 | 5 | 38.4 |
| *Abatus cordatus* | - | - | - | - | 497 | 2 | 36.8 |
| *Acanthaster planci* (OG) | - | - | - | - | 864 | 8 | 63.6 |
| *Amphiura filiformis* (OG) | 180.1 | 61,558,173 | 52,206,727 | 416,946 | 761 | 8 | 55.9 |
| *Apatopygus recens* | 415.3 | 74,315,687 | 56,898,850 | 274,380 | 1152 | 5 | 85.2 |
| *Apostichopus japonicus* (OG) | - | - | - | - | 849 | 9 | 62.4 |
| *Araeosoma leptaleum* | - | - | - | - | 1184 | 2 | 87.8 |
| *Arbacia lixula* | - | - | - | - | 1234 | 1 | 91.6 |
| *Aspidodiadema hawaiiense* | 228.7 | 109,716,219 | 104,518,714 | 311,032 | 1060 | 11 | 77.9 |
| *Asterias rubens* (OG) | 230.4 | 31,890,613 | 25,495,009 | 103,090 | 805 | 9 | 59.1 |
| *Asthenosoma varium* | - | - | - | - | 1254 | 7 | 92.6 |
| *Astrophyton muricatum* (OG) | 195.9 | 25,191,954 | 22,281,829 | 149,146 | 478 | 5 | 35.1 |
| *Bathysalenia phoinissa* | - | - | - | 154,120 | 605 | 6 | 44.5 |
| *Brissus obesus* | - | - | - | - | 773 | 0 | 57.4 |
| *Caenopedina hawaiiensis* | - | - | - | - | 1094 | 2 | 81.1 |
| *Clypeaster japonicus* | 198.6 | 10,505,520 | 9,298,316 | 74,743 | 829 | 1 | 61.5 |
| *Clypeaster rosaceus* | - | - | - | - | 1242 | 2 | 92.1 |
| *Clypeaster subdepressus* | - | - | - | - | 1233 | 3 | 91.4 |
| *Colobocentrotus atratus* | - | - | - | - | 1158 | 0 | 86.0 |
| *Conolampas sigsbei* | - | - | - | - | 1001 | 5 | 74.0 |
| *Cystechinus* c.f. *giganteus* | - | - | - | - | 661 | 0 | 49.1 |
| *Dendraster excentricus* | - | - | - | - | 337 | 1 | 25.0 |
| *Diadema setosum* | - | - | - | - | 305 | 1 | 22.6 |
| *Echinarachnius parma* | - | - | - | - | 1187 | 3 | 88.0 |
| *Echinocardium cordatum* | - | - | - | - | 1012 | 2 | 75.0 |
| *Echinocardium mediterraneum* | - | - | - | - | 1185 | 1 | 88.0 |
| *Echinocyamus crispus* | - | - | - | - | 709 | 6 | 52.2 |
| *Echinometra mathaei* | - | - | - | - | 1055 | 0 | 78.4 |
| *Echinothrix calamaris* | 203.9 | 68,871,087 | 60,443,904 | 251,788 | 1252 | 4 | 92.7 |
| *Encope emarginata* | 206.2 | 12,015,888 | 10,461,870 | 66,241 | 1076 | 1 | 79.9 |
| *Eucidaris metularia* | 321.6 | 38,802,159 | 29,047,401 | 83,318 | 412 | 2 | 30.5 |
| *Eucidaris tribuloides* | - | - | - | - | 414 | 0 | 30.8 |
| *Evechinus chloroticus* | - | - | - | - | 1196 | 1 | 88.8 |
| *Fellaster zelandiae* | 313.9 | 62,619,791 | 51,742,748 | 168,598 | 1269 | 1 | 94.2 |
| *Heliocidaris erythrogramma* | - | - | - | - | 931 | 0 | 69.2 |
| *Histocidaris variabilis* | 329.2 | 84,103,672 | 73,884,180 | 144,044 | 1189 | 5 | 88.0 |
| *Holothuria forskali* (OG) | - | - | - | - | 743 | 8 | 54.6 |
| *Lamberticrinus messingi* (OG) | 195.0 | 22,989,820 | 20,234,597 | 59,049 | 351 | 3 | 25.9 |
| *Leodia sexiesperforata* | 203.8 | 16,211,182 | 14,174,475 | 68,964 | 1064 | 1 | 79.0 |
| *Leptosynapta clarki* (OG) | - | - | - | - | 403 | 4 | 29.6 |
| *Lissodiadema lorioli* | - | - | - | - | 770 | 0 | 57.2 |
| *Loxechinus albus* | - | - | - | - | 1265 | 1 | 93.9 |
| *Lytechinus variegatus* | - | - | - | - | 1257 | 2 | 93.2 |
| *Mellita tenuis* | - | - | - | - | 1002 | 1 | 74.4 |
| *Meoma ventricosa* | - | - | - | - | 1053 | 0 | 78.2 |
| *Mesocentrotus nudus* | - | - | - | - | 1247 | 0 | 92.6 |
| *Metacrinus rotundus* (OG) | 198.5 | 23,791,832 | 20,900,345 | 83,718 | 642 | 7 | 47.2 |
| *Micropyga tuberculata* | - | - | - | 170,514 | 415 | 4 | 30.5 |
| *Ophiocoma echinata* (OG) | 278.5 | 28,427,026 | 24,836,025 | 130,153 | 712 | 7 | 52.4 |
| *Paracentrotus lividus* | - | - | - | - | 1266 | 0 | 94.1 |
| *Patiria miniata* (OG) | - | - | - | - | 740 | 8 | 54.4 |
| *Peronella japonica* | 208.3 | 17,696,287 | 15,707,931 | 106,110 | 1043 | 6 | 77.0 |
| *Prionocidaris baculosa* | - | - | - | - | 794 | 3 | 58.8 |
| *Psammechinus miliaris* | - | - | - | - | 1173 | 1 | 87.1 |
| *Rhyncholampas pacificus* | 346.8 | 63,116,413 | 52,160,741 | 234,910 | 1070 | 2 | 79.3 |
| *Saccoglossus kowalevskii*<br>(OG) | - | - | - | - | 668 | 6 | 49.2 |
| *Scaphechinus mirabilis* | 401.7 | 10,169,195 | 8,796,067 | 127,664 | 974 | 4 | 72.1 |
| *Sinaechinocyamus mai* | 297.2 | 60,208,172 | 52,265,201 | 164,646 | 1233 | 1 | 91.5 |
| *Sphaerechinus granularis* | - | - | - | - | 1214 | 1 | 90.1 |
| *Sterechinus neumayeri* | 207.3 | 26,376,279 | 21,308,840 | 122,001 | 1097 | 1 | 81.4 |
| *Stereocidaris nascaensis* | 327.0 | 69,962,495 | 60,316,050 | 121,508 | 1200 | 6 | 88.7 |
| *Stomopneustes variolaris* | - | - | - | - | 908 | 0 | 67.5 |
| *Strongylocentrotus*<br>*purpuratus* | - | - | - | - | 1266 | 5 | 93.7 |
| *Tetrapygyus niger* | 202.5 | 63,040,541 | 52,761,671 | 163,084 | 1287 | 2 | 95.5 |
| *Tripneustes gratilla* | 154.6 | 62,962,239 | 57,015,692 | 130,376 | 1309 | 1 | 97.2 |
| *Tromikosoma hispidum* | 300.4 | 57,208,357 | 47,909,038 | 234,502 | 1238 | 7 | 91.5 |

